# Pentapeptide repeat proteins QnrB1 and AlbG require ATP hydrolysis to rejuvenate poisoned gyrase complexes

**DOI:** 10.1101/2020.09.24.310243

**Authors:** Łukasz Mazurek, Dmitry Ghilarov, Elizabeth Michalczyk, Zuzanna Pakosz, Wojciech Czyszczoń, Karolina Wawro, Iraj Behroz, Roderich D. Süssmuth, Jonathan G. Heddle

## Abstract

DNA gyrase, a type II topoisomerase found predominantly in bacteria, is the target for a variety of “poisons”, namely natural product toxins (e.g. albicidin. microcin B17) and clinically important synthetic molecules (e.g. fluoroquinolones). Resistance to both groups can be mediated by pentapeptide repeat proteins (PRPs). Despite long-term studies, the mechanism of action of these protective PRPs is not known. We compared activities of two such proteins, QnrB1 and AlbG *in vitro*. Each of them provided specific protection against its cognate toxin (fluoroquinolone or albicidin), which strictly required ATP hydrolysis by gyrase. Through a combination of fluorescence anisotropy, pull-downs and photocrosslinking we show that QnrB1 binds to the GyrB protein. We further probed the QnrB1 binding site using site-specific incorporation of a photoreactive amino acid and mapped strong and specific crosslinks to the N-terminal ATPase/transducer domain. We propose a model in which protective PRPs bind to the enzyme as T-segment DNA mimics to promote dissociation of the bound poison molecule.

## INTRODUCTION

DNA gyrase is a type II topoisomerase found predominantly in bacteria, but also in archaea, plant chloroplasts and *Apicomplexa* (1–4). Gyrase is an essential enzyme responsible for the maintenance of the bacterial chromosome in a negatively supercoiled state and removal of torsion accumulated in front of DNA and RNA polymerases, allowing replication and transcription to proceed (5, 6). Gyrase functions as a heterotetramer consisting of two GyrA and two GyrB subunits, with three interfaces (“gates”) between them (**Figure 1A**). The supercoiling mechanism of gyrase has been extensively studied and is understood in some detail (1, 4, 5) (**Figure 1A**). Briefly, double-stranded DNA is wrapped around the C-terminal domains of GyrA (CTD) such that one DNA segment (the gate segment, G) is bound across the “DNA gate” (formed by the N-terminal winged helix domains (WHD) of GyrA and the C-terminal topoisomerase-primase (TOPRIM) domains of GyrB) whilst a more distal segment of the same DNA molecule (T, or transported segment) forms a positive node with the G segment. In the presence of Mg^2+^ ions, DNA is cleaved by the two active-centre Tyr residues of the GyrAs, each of which forms a new phosphodiester bond with the cleaved fragment. ATP binding to the N-terminal ATPase domains of GyrB induces their dimerization. This leads to the capture of the T-segment and its passage through the cleaved G segment and out of the bottom gate of the enzyme (**Figure 1A**).

**Figure 1.**
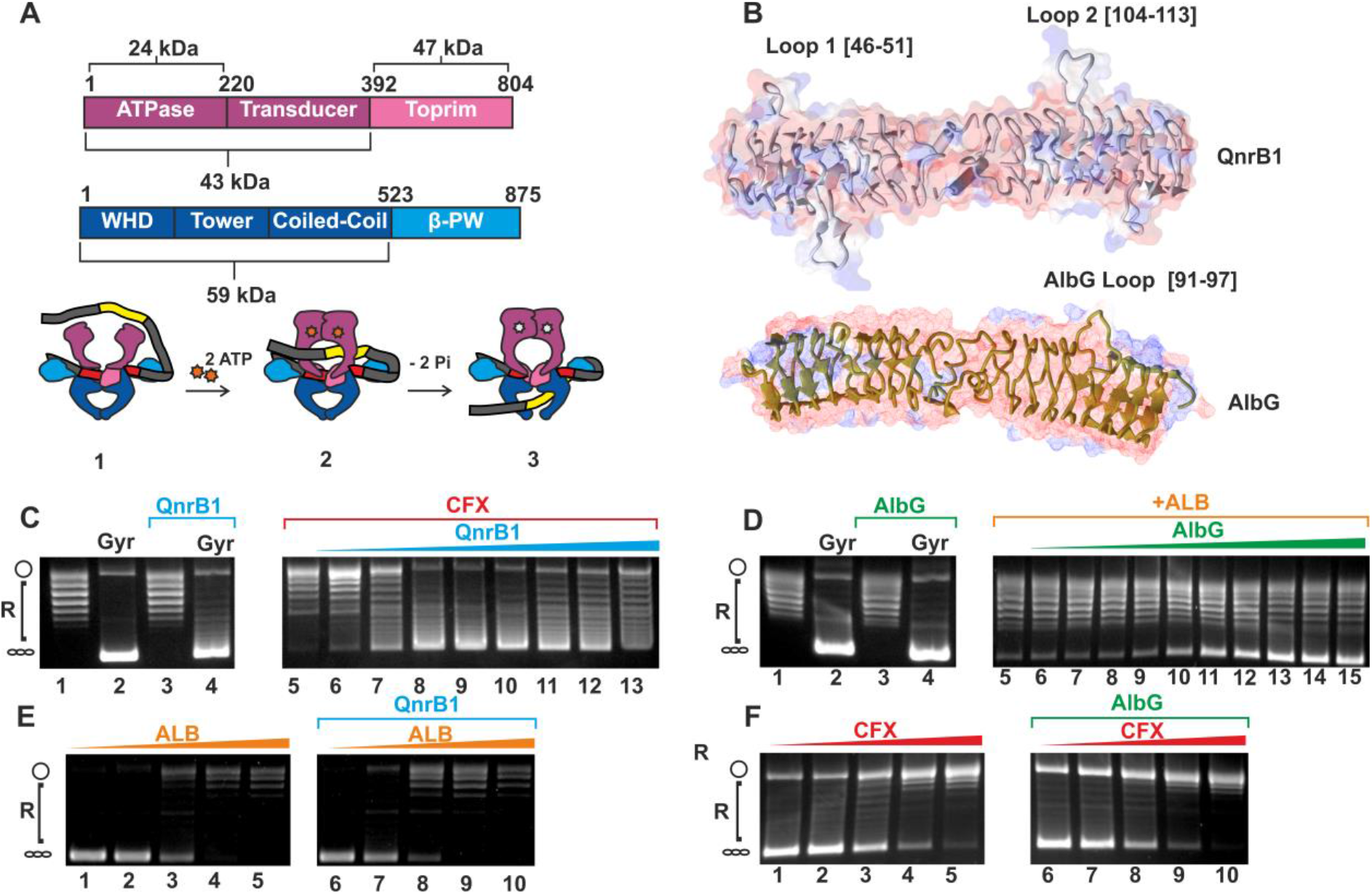
Poison-specific gyrase rescue mediated by QnrB1 and AlbG proteins. (**A**) *Top:* Domain architecture of *E. coli* DNA gyrase. Gyrase B is composed of ATPase, transducer (both purple) and TOPRIM (pink) domains. Indicated are molecular masses of different protein fragments (24 kDa, 43 kDa and 47 kDa) produced and used in this work. Gyrase A is composed of winged-helix (WHD), tower, coiled-coil (all three blue) and β – pinwheel (cyan) domains. *Below*: A scheme of DNA gyrase mechanism. (**1**) A double-stranded DNA (G-segment (red)) is captured by the DNA gate, the ATPase gate is open to allow enter of the T-segment (yellow); (**2**) ATP binding leads to dimerization of the ATPase domains and trapping of the T-segment in the upper cavity; (**3**) this is followed by ATP hydrolysis, G-segment cleavage, DNA-gate opening and T-segment translocation. (**B**) Cartoon and molecular surface (transparent) representations of two PRPs used in this study – QnrB1 PDB: 2XTW (*top*) and AlbG PDB: 2XT2 (*bottom*). Indicated are looped regions, previously found to be implicated in protective activity. (**C**). Plasmid supercoiling assay showing: (*left*) lane 1: relaxed pBR322, lane 2: supercoiling by 1 U of gyrase, lane 3: lack of detectable nuclease activity in the purified QnrB1 (50 μM QnrB1) and lane 4: partial inhibition of gyrase by 50 μM QnrB1; (*right*) lanes 6-13: gyrase inhibition by CFX (5 μM) and rescue by increasing concentrations of QnrB1 (0.04; 0.2; 1; 5; 10; 20; 25; 50 μM). Positions of negatively supercoiled and relaxed DNA are indicated by the graphics at the left (same notation used in other figures). (**D**) Plasmid supercoiling assay showing (*left*): lane 1: relaxed pBR322; lane 2: supercoiling by 1 U of gyrase; lane 3: lack of detectable nuclease activity in the purified AlbG (50 μM) and lane 4: lack of effect of 50 μM AlbG on gyrase supercoiling; (*right*) gyrase inhibition by ALB (3.3 μM) and partial rescue by increasing concentrations of AlbG (0.0016; 0.008; 0.04; 0.2; 1; 5; 10; 20; 25; 50 μM) (**E**). *Left* gel shows gyrase supercoiling inhibition by increasing concentrations of albicidin lanes 1-5 (0.016; 0.16; 1.6; 16; 160 [μM]). *Right* gel, effect of the addition of 5 μM QnrB1 to the same reactions (lanes 6 – 10) (**F**). *Left*, lanes 1-5: gyrase supercoiling inhibition by increasing concentrations of ciprofloxacin (0.016; 0.08; 0.4; 2; 10 μM); *right*, lanes 6 – 10: effect of adding 5 μM AlbG to the reactions.

The most effective gyrase-targeting agents are the so-called “poisons”, which trap the enzyme in a post G-segment cleaved state. Accumulation of these stalled enzyme complexes on DNA leads to cell death (7). The clinically important fluoroquinolone (FQs) are a particularly successful class of antibiotics, having a broad-range bactericidal activity. FQs intercalate between the DNA bases and are anchored to the Ser83 (*Escherichia coli* numbering) and Glu87 residues of GyrA via a Mg^2+^ ion (8). Additional interactions involve GyrB residues located near the FQ C-7 ring. While mutations in GyrA account for the majority of target-related resistance to FQs, concerns have grown in recent years over the spread of plasmid-borne resistance, which is in part mediated by *qnr* (quinolone resistance) genes (9). Although resistance conferred by *qnr* genes is incomplete, it facilitates the emergence of chromosomal *gyrA/B* mutations (10). *Qnr* family members encode pentapeptide repeat proteins (PRPs), composed from tandem 5-amino acid repeats of general sequence A(D/N)LXX) (11). A homologous protein, MfpA, found in *Mycobacterium tuberculosis* increases resistance of this organism to fluoroquinolones (12). Likewise, AlbG from *Xanthomonas albilineans* and McbG from *E. coli* (13, 14) are thought to protect the host gyrase against specific gyrase-targeting natural products. *AlbG* is located within a cluster of genes responsible for the synthesis of a polyketide antibiotic albicidin (ALB) (15). Albicidin interacts with GyrA and leads to the accumulation of cleaved DNA-enzyme complexes, however, details of the mechanism and mode of binding are not known (15). *McbG* is a part of the *E. coli* microcin B17 (MccB17) biosynthetic cluster. MccB17 is a 3 kDa post-translationally modified peptide, decorated by thiazole and oxazole heterocycles (16); similarly to albicidin and FQs, it stabilises the gyrase cleavage complex (17, 18). Partial cross-resistance with quinolones was also reported for both albicidin MccB17 (15, 19), suggesting that the binding sites for the compounds may overlap.

Structures of AlbG, MfpA, QnrB1 and other topoisomerase-interacting PRPs have been solved by x-ray crystallography (13, 20–23). All these proteins fold into rod-like β-barrels with a square cross-section and dimerise via a C-terminal α-helix (**Figure 1B**). Structure-functional analysis of QnrB1 highlighted the significance of loops protruding from the helical scaffold (loops 1 and 2) as their deletion abrogated protective activity (21, 23, 24). Various hypotheses have been proposed to explain the protective effects of QnrB1 and related PRPs. The right-helical shape and stretches of negative charge along the length of MfpA and other PRPs are similar to that of double-stranded DNA, leading to the idea that PRPs can act as G-segment mimics (20). In this concept, as a result of competition between the PRP and G-segments for gyrase binding, there would be fewer complexes with DNA present, hence fewer lethal double stranded DNA breaks. In agreement with this hypothesis, MfpA was shown to inhibit gyrase *in vitro* (20, 25) and Qnr protein was shown to reduce DNA binding to gyrase in a filter assay (26). However, later it was shown that QnrB1 does not inhibit gyrase supercoiling activity, at least at the concentrations required for rescue (21, 22, 25). This observation is incompatible with a model in which PRPs compete with G-segment binding, as clearly supercoiling cannot proceed without G-segment present. This led to the idea of a direct recognition of the gyrase-drug complex by the PRP, which results in loss of the drug from gyrase (21). These observations and apparent DNA-mimicry can be reconciled by the proposed “T-segment mimicry” model, where the transport of the PRP through the enzyme destabilises the enzyme-drug complex and allows for dissociation of the drug from the enzyme (27).

In order to clarify the mechanisms whereby PRPs protect gyrase from poisons, we have analysed the protective effects of QnrB1, AlbG and McbG towards their cognate target toxins (CFX, ALB or MccB17) *in vivo* and *in vitro* using *E. coli* DNA gyrase as a model. We have shown that the protective effect in each case is specific and generally does not lead to the inhibition of gyrase. The rescue effect strictly requires ATP hydrolysis by gyrase and leads to the destabilisation of the enzyme-drug complex. Our data are not compatible with the G-segment displacement model but agree with the models where interactions of PRP with gyrase are transient and drug-specific. Correlating results have been independently obtained by Dr. Lipeng Feng, Dr. Anthony Maxwell and Dr. David Lawson for *Mycobacterium smegmatis* MfpA protein (personal communication).

## MATERIALS AND METHODS

### Bacterial strains, plasmids and molecular cloning

*Escherichia coli* DY330 GyrA-SPA (sequential peptide affinity) (W3110 Δ*lacU169 gal490 λcI857* Δ(cro-bioA) *gyrA*-SPA) and GyrB-SPA (W3110 Δ*lacU169 gal490 λcI857* Δ(cro-bioA) *gyrB*-SPA) (28) and BW25113 were gifts from Dmitry Sutormin (Skolkovo Institute of Science and Technology).

Plasmids pET28-*albG* (encoding AlbG with a thrombin-cleavable His-tag, (13), pBAD-*mcbABCDEFG* (*wt* MccB17 operon) and pET28-*qnrB1* (encoding QnrB1 with a C-terminal 6xHis tag) are gifts from Dr. Mikhail Metelev (Uppsala University). Plasmids pET21-GyrA and pET21-GyrB coding for full-length untagged GyrA and GyrB subunits amplified from E. coli MG1655 by PCR and cloned between NdeI and BamHI sites were previously produced in the laboratory. Plasmids pET21-3xFLAG-GyrB and pET21-GyrA-FLAG were constructed by amplifying GyrA and GyrB genes (as above) with N-terminal (GyrB) 3xFLAG tag and C-terminal (GyrA) FLAG tags and cloning using Nde I and Xho I sites into pET21. pET28-GyrB47 was constructed by amplifying coding sequence for gyrase TOPRIM domain (393-804) from *E. coli* MG1655 and cloning it into pET28 using Nde I and Xho I restriction sites behind the N-terminal hexahistidine tag. pAJR10.18 (GyrA59) and pAJ1 (GyrB43) were gifts of A. Maxwell, John Innes Centre. For pBAD-*albG*, pBAD-*mcbG* and pBAD-*qnrB1*, corresponding genes were amplified from the corresponding template plasmids and cloned into pBAD vectors using Nco I and Xho I restriction sites.

### Purification of gyrase subunits and domains

*E. coli* full-length GyrA and GyrB subunits were purified similarly to (29). Plasmids pET21b containing corresponding genes were transformed into BL21 (DE3) Gold (Agilent). For GyrA, 2 L of TB culture supplemented with 100 μg/ml ampicillin was incubated at 37°C with shaking to OD_600_ = 0.75, induced with 500 μM isopropyl-β-d-thiogalactopyranoside (IPTG) and incubated for a further 4 hours at 37°C. Cells were harvested by centrifugation at 6000 g for 15 minutes at 4°C. Pellets were resuspended in buffer A (50 mM Tris-HCl pH 7.5, 10% glycerol, 1 mM EDTA, 2 mM DTT) and supplemented with protease inhibitor cocktail (Pierce). Cells were lysed using a French press and cell debris removed by centrifugation at 40000 g for 20 minutes at 4°C. Clarified lysate was loaded onto 10 ml Q XL column (GE Healthcare) in buffer A, washed with 10 CV of buffer A, 10 CV of buffer A supplemented with 0.1 M NaCl and eluted with gradient (0.1-1 M) of NaCl in buffer A over 5 CVs. Collected fractions were dialysed into buffer A overnight at 4°C and further purified by MonoQ HR 16/10 (20 ml) using the same method as described for Q XL. Peak fractions were pooled together, concentrated using Amicon concentrators (Millipore) and loaded onto a Superdex 200 Increase 10/300 (Cytiva) column equilibrated in buffer A. Fractions containing GyrA were aliquoted, frozen in liquid nitrogen and stored at −80°C in buffer A. For GyrB, 2 L of TB culture was grown at 37°C with shaking to OD_600_ = 0.8, induced with 500 μM IPTG and incubated for a further 3 hours at 28°C. Cells were lysed and processed in buffer A as described for GyrA and protein was purified using heparin affinity (Heparin FF 16/10 column) and anion exchange (MonoQ HR 16/10 column) chromatography (Cytiva), eluting with a 0-1 M gradient of NaCl. Peak fractions from MonoQ were pooled, dialysed into buffer A, aliquoted and frozen at −80°C. FLAG-tagged versions of both GyrA and GyrB were purified similarly.

GyrB 43 kDa domain (GyrB43) was purified similarly to (30). The plasmid pAJ1(31) was transformed into BL21 (DE3) Gold cells. 1 L culture of TB inoculated with transformed cells was incubated at 37°C with shaking to OD_600_ = 0.8 after which it was induced with 500 μM IPTG. The temperature was reduced to 25°C and the culture was incubated with shaking overnight. Cells were harvested and processed similarly to GyrA and GyrB proteins. GyrB43 was first purified on Q XL similarly to GyrA and B proteins. Fractions containing protein of interest were combined and ammonium sulfate was added to 1.5M. Salt-adjusted protein was loaded to 10 ml Phenyl Sepharose HS FF (Cytiva) column equilibrated in 1.5M (NH_4_)_2_SO_4_ and protein was eluted by 20 CV gradient from 1.5 M to 0 (NH_4_)_2_SO_4_. Collected fractions were pooled together, dialysed overnight in buffer A and loaded onto a 20 ml MonoQ column as described for GyrA and GyrB proteins. Peak fractions from MonoQ were pooled, dialysed overnight into buffer A, concentrated to 10 mg/ml, aliquoted, frozen in liquid N_2_ and stored at −80°C.

GyrB47 purification was described in (32). 2 L culture of BL21 (DE3) Gold pET28-GyrB47 was grown in TB at 37°C. At OD_600_ = 0.75 IPTG was added to 0.5 mM and cells were grown for further 4 hours at 37°C. After harvesting, cells were resuspended in a lysis buffer (50 mM Tris-Cl pH 7.5, 150 mM NaCl, 20 mM imidazole, 10% glycerol) and lysed using a French press. Cleared lysate was loaded on a pre-equilibrated 5 ml HisTrap FF column (Cytiva), washed with the lysis buffer and eluted with the lysis buffer containing 250 mM imidazole. Peak fractions were dialysed overnight into buffer A and further purified on a 5 ml Q HP column (Cytiva). A step gradient of NaCl was used and protein eluted at 40% NaCl.

GyrA59 was purified similarly to (33). Plasmid pAJR10.18 was transformed into BL21(DE3) Gold cells. 2 L of culture was grown in LB at 37°C to OD_600_ = 0.85, induced with 0.5 mM IPTG, and grown for further 4 hours at 37°C. Cells were lysed by sonication in buffer A. Lysate was cleared by centrifugation at 87 000 g for 30 minutes at 4°C and GyrA59 was purified first on 16/10 Heparin FF (Cytiva) column using 0.1-1 M gradient of NaCl in buffer A. After overnight dialysis to buffer A, GyrA59 was further purified on MonoQ (16/10, Cytiva) using a gradient 0-0.7M NaCl over 6 CV. Fractions containing GyrA59 were pooled, concentrated and loaded onto Superdex 200 16/600 gel filtration column. Pure protein, concentrated to ~5 mg/ml was aliquoted, flash-frozen in liquid N_2_ and stored at −80°C.

### AlbG purification

Plasmid pET28-AlbG containing the *albG* open reading frame with a thrombin-cleavable N-terminal hexahistidine tag was transformed into BL21 (DE3) Gold cells. 3 L of 2xYT liquid media supplemented with 30 μg/ml kanamycin were inoculated using 1:100 ratio of starter overnight culture and incubated at 37°C with shaking to OD_600_ = 0.8. After induction with 500 μM IPTG the temperature was reduced to 24°C and the cultures were incubated overnight with shaking. Cells were harvested by centrifugation at 7000 g for 30 minutes at 4°C. Pellets were resuspended in AlbG lysis buffer (50 mM Tris-Cl pH 8, 20 mM imidazole, 300 mM NaCl, 5% glycerol) supplemented with protease inhibitors (Pierce). Resuspended cells were incubated for 30 minutes on ice with 2 mg/ml lysozyme and 5 μg/ml DNase I. Cells were lysed using a French press and lysate was cleared by centrifugation at 87 000 g for 30 minutes at 4°C. AlbG was purified by Ni-affinity chromatography (5ml HisTrap HP, Cytiva), dialysed overnight into buffer A and further purified by anion exchange (Q XL 5 ml, Cytiva) using step elution by increasing concentrations of NaCl. Peak fractions eluted between 0.2-0.3 M NaCl were dialysed into buffer A, concentrated and stored at −80 C.

### QnrB1 purification

Plasmid pET28-QnrB1 containing the *qnrB1* open reading frame with C-terminal hexa-histidine tag was transformed into BL21 (DE3) Gold cells 3 L of TB liquid media supplemented with 30 μg/ml kanamycin were inoculated with1/100 ratio of starter overnight culture and incubated at 37°C with shaking to OD_600_ = 0.8. After induction with 500 μM IPTG the temperature was reduced to 24°C and the cultures were incubated overnight with shaking. Cells were harvested by centrifugation at 7000 g for 30 minutes at 4°C. Pellets were resuspended in QnrB1 lysis buffer (50 mM Tris-Cl pH 8.0, 200 mM (NH_4_)_2_SO_4_, 10% glycerol, 20 mM imidazole) supplemented with protease inhibitors (Pierce). Resuspended cells were incubated for 30 minutes on ice with 1 mg/ml lysozyme with periodic mixing. Cells were lysed using a French press and lysate was cleared by centrifugation at 87 000 g for 30 minutes at 4°C. QnrB1 was purified by Ni-affinity chromatography (HisTrap HP 5 ml, Cytiva). The column was equilibrated with QnrB1 lysis buffer and after loading the lysate washed with 20 CVs of wash buffer (50 mM Tris-Cl pH 8.0, 200 mM (NH_4_)_2_ SO_4_, 10% glycerol, 50 mM imidazole). The fractions were eluted with an elution buffer (50 mM Tris-Cl pH 8.0, 200 mM (NH_4_)_2_SO_4_, 10% glycerol, 250 mM imidazole). Fractions containing protein were pooled and dialysed overnight against buffer A-Arg (50 mM Tris-HCl pH 7.5, 50 mM arginine hydrochloride, 10% glycerol, 1 mM EDTA, 2 mM DTT) and loaded on MonoQ HR 16/10 ion exchange column (Cytiva). Peak fractions eluted by 0-1 M NaCl gradient were pooled, concentrated, and loaded onto a 10/300 Superdex S75 Increase SEC column (Cytiva) previously equilibrated with QnrB1 storage buffer (20 mM Tris pH 7.5, 50 mM NaCl, 5% glycerol, 50 mM arginine hydrochloride, 2 mM DTT). Peak fractions were concentrated, snap-frozen in liquid N_2_ and stored at −80°C.

### Purification of QnrB1 *para*-benzoyl-phenylalanine (*p*Bpa) mutants

Plasmids pBAD-*qnrB1[x]pBpa*, carrying *qnrB1* amber mutants with N-terminal 6xHis, were transformed into One Shot BL21 Star (DE3) cells (Thermo) together with the pEVOL-*p*Bpa plasmid (34). 100 ml of LB liquid media supplemented with 30 μg/ml kanamycin and 100 μg/ml ampicillin was inoculated using a 1:100 ratio of starter overnight culture and incubated at 37°C with shaking to OD_600_ = 0.3. At this point *para*-benzoyl-phenylalanine (pBpa) was added to a final concentration of 1 mM. The incubation was continued until the culture reached OD_600_ = 0.6. The expression of orthogonal aaRS was induced by the addition of arabinose (10 mM). After 6 h of expression at 37°C with shaking, the cells were collected by centrifugation at 7000 g for 30 minutes at 4°C. Pellets were resuspended in QnrB1 lysis buffer (50mM Tris-Cl pH 8.0, 200 mM (NH_4_)_2_SO_4_, 10% glycerol, 20 mM imidazole) supplemented with protease inhibitors (Pierce). Resuspended cells were incubated with mixing for 30 minutes on ice with 1 mg/ml lysozyme. Cells were lysed by sonication and lysate was cleared by centrifugation at 87 000 g for 30 minutes at 4°C. Lysate was loaded onto a Ni^2+^-affinity chromatography column (5 ml HisTrap HP, Cytiva) equilibrated with QnrB1 lysis buffer. After loading the lysate, the column was washed with 20 column volumes of wash buffer (50 mM Tris-Cl pH 8.0, 200 mM (NH_4_)_2_SO_4_, 10% glycerol, 50 mM imidazole). The fractions were eluted with the elution buffer (50 mM Tris-Cl pH 8.0, 200 mM (NH_4_)_2_SO_4_, 10% glycerol, 250 mM imidazole). Fractions containing protein were subsequently diluted with the storage buffer (20 mM Tris-Cl pH 7.5, 50 mM NaCl, 5% glycerol, 50 mM arginine hydrochloride), concentrated to desired concentration and stored at −80°C.

### Purification of gyrase-targeting toxins

Microcin B17 was obtained from *Escherichia coli* BW25113 transformed with *pBAD-mcbABCDEFG* plasmid using a method described previously (35). Briefly, transformants were cultured in 2 L of 2xYT media until OD_600_ = 0.7, induced with 10 mM arabinose and left overnight at 37°C. Cells were collected by centrifugation at 4000 g, resuspended in 100 mM acetic acid/1 mM EDTA solution and lysed by boiling for 10 min on a water bath. Lysate was clarified by centrifugation (12 000g) and loaded onto Agilent BondElute 10 g C18 cartridge, equilibrated with 0.1% TFA. Cartridge was extensively washed first with 20 CV 0.1% TFA, then with 20 CV of 10% MeCN in 0.1% TFA before elution with 30% MeCN/0.1% TFA. Eluate was dried *in vacuo*, reconstituted in 10% DMSO and further purified by HPLC using COSMOSIL 5C18-MS-II 120Å 5μm, 10.0x 150 mm column and 10-30% gradient of MeCN. Albicidin was produced by total synthesis (36).

### MIC measurements

Minimal inhibitory concentrations of compounds were measured by broth microdilution in 96-well plates as described by the Clinical & Laboratory Standards Institute (37). All measurements were performed as triplicates. The error is expressed by standard deviation of the mean.

### Gyrase activity assays

For supercoiling assays, 1 unit of *E.coli* gyrase (1 unit defined as the amount of gyrase required for full conversion of 500 ng relaxed pBR322 DNA into the completely supercoiled form in 30 min at 37°C in a 30 μl reaction, corresponded to 4 nM concentration) was incubated in a total volume of 30 μl in assay buffer (35 mM Tris-Cl pH 7.5, 24 mM KCl, 4 mM MgCl_2_, 2 mM DTT, 1.8 mM spermidine, 1 mM ATP, 6.5 % (w/v) glycerol, 0.1 mg/ml albumin) and 500 ng relaxed pBR322 DNA (Inspiralis Ltd.). The compounds and/or proteins were replaced by corresponding buffers in control reactions. Reactions were stopped by the addition of chloroform:isoamyl alcohol (24:1) and STEB (20% sucrose, 50 mM Tris-Cl pH 8, 5 mM EDTA, 0.25 mg/ml bromophenol blue). The aqueous layers from the assays were run on 1% agarose TAE gels at 80 V for 2.5 hours in 1 x TAE buffer. Once complete, the gels were stained with 10 μg/ml ethidium bromide solution (Sigma) for 15 minutes and de-stained with 1 x TAE buffer for 15 minutes and visualised using a gel documentation system (UVP). Quantification of the amount of supercoiled DNA was performed using Fiji software (38). Percentage of supercoiled DNA was plotted against QnrB1 concentration and the data was fitted to the equation: %*SC* = *ae*^*b[QnrB1]*^, where a and b are function parameters found after fitting the function using Origin (Pro), Version 2020 b. IC_50_ (concentration of QnrB1 required for 50% supercoiling inhibition). Gyrase relaxation assays were carried out in a similar manner, but ATP and spermidine were omitted and ~5 U of gyrase were used. For gyrase cleavage assays ~5 U of gyrase were used and reactions were terminated by the addition of 0.2% sodium dodecyl sulphate (SDS) and 0.2 mg/ml proteinase K, followed by incubation for 30 minutes at 37°C prior to chloroform extraction and gel analysis. For cleavage of short DNA fragments, 20 nM fragment (with the exception of the 76 bp fragment where 30 nM was used as indicated in figure) was incubated with 5U of gyrase for 30 min, followed by reaction termination as described above. Reaction products were separated on 4-20% TBE polyacrylamide gels (Thermo). DNA fragments (300-100 bp) were obtained by amplification from the pBR322 template, following by purification (GeneJet Gel Extraction and DNA Cleanup Micro kit, Thermo). 76 bp fragment was ordered as a pair of complementary oligos and annealed in a PCR machine.

Time courses of gyrase-mediated DNA cleavage were performed as follows: *E.coli* gyrase (100 units) was incubated at 25°C in 400 μl reactions with assay buffer (35 mM Tris-Cl pH 7.5, 24 mM KCl, 4 mM MgCl_2_, 2 mM DTT, 1.8 mM spermidine, 6.5 % (w/v) glycerol, 0.1 mg/ml albumin) and 500 ng relaxed pBR322 DNA (Inspiralis Ltd). For Ca^2+^ induced cleavage, MgCl_2_ was replaced with CaCl_2_. The buffer was supplemented with 1 mM ATP or 0.5 mM ADPNP as required. When tested compounds and/or proteins were absent in control reactions, they were replaced by DMSO or corresponding buffers. At selected time points, 20 μl aliquots were withdrawn and stopped by addition of 2 μl 5% SDS and 2 μl 250 mM EDTA. After the time course, collected samples were treated with proteinase K (0.2 mg/ml) for 30 minutes at 37°C and extracted by chloroform:isoamyl alcohol (24:1). The aqueous layers from the assays were mixed with STEB and run on 1% agarose TAE gels at 80 V for 2.5 hours in 1 x TAE buffer. Once complete, the gels were stained with 10 μg/ml ethidium bromide solution (Sigma) for 15 minutes and de-stained with 1xTAE buffer for 15 minutes and visualised using a gel documentation system (UVP).

Cleavage complex stability assays were performed as follows: an initial 60 μl reaction was set up using 80 units of gyrase in the assay buffer, 50 nM of relaxed DNA and 20 μM CFX. which was incubated at 37°C for 10 minutes. Then 20 ul aliquots were withdrawn and diluted 20-fold with the buffer supplemented with 5 μM QnrB1/ 5 μM AlbG and/or 0.5 mM ATP as required. For AlbG containing assays 1 μM of albicidin was used to form a starting complex. At chosen time points 20 μl from each reaction was pipetted to a separate tube and stopped by adding 2 μl of 5% SDS and 2 μl 250 mM EDTA. After the time course, samples were treated with proteinase K (0.2 mg/ml) for 30 minutes at 37°C, then a chloroform-isoamyl alcohol mixture (24:1) and STEB (20% sucrose, 50 mM Tris-Cl pH 8, 5 mM EDTA, 0.25 mg/ml bromophenol blue) were added. The aqueous layers from the assays were run on 1% agarose TAE gels at 80 V for 2.5 hours in 1 x TAE buffer. Once complete, the gels were stained with 10 μg/ml ethidium bromide solution (Sigma) for 15 minutes and de-stained with 1 x TAE buffer for 15 minutes and visualised using a gel documentation system (UVP).

### Electrophoretic mobility shift assays (EMSA)

147 bp pBR322 dsDNA fragment with known strong gyrase binding site (39) was produced by PCR and purified with Thermo Scientific GeneJet Gel Extraction and DNA Cleanup Micro kit. 20 nM of the fragment was mixed with 0.2 μM of reconstituted gyrase in EMSA buffer (30 mM Tris-Cl pH 7.5, 75 mM KCl, 6% glycerol, 2 mM MgCl_2_, 1 mM DTT). QnrB1 or AlbG were added as indicated. In control reactions, QnrB1/AlbG were supplemented by their storage buffers added. Reactions were incubated for 30 min at 25°C and run on 6% polyacrylamide gels in TBM buffer (90 mM Tris-borate, pH 7.5, 4 mM MgCl_2_) at 150V at room temperature. After the run gels were stained with SYBR Gold (Thermo) for 20 mins and visualized under UV light.

### ATPase assays

Gyrase and GyrB 43 ATPase assays were carried out using Inspiralis kits according to the protocol provided by Inspiralis Ltd based on Tamura & Gellert (40). Each reaction contained 50 mM Tris.HCl (pH 7.5), 1 mM EDTA, 5 mM magnesium chloride, 5 mM DTT, 10 % (w/v) glycerol, 0.8 mM PEP, 0.4 mM NADH and ~1U of PK/LDH mix (Sigma). For GyrB43 assays, the concentration of GyrB43 was 4 μ*M*. For gyrase assays, 50 nM gyrase tetramer (A2B2) was used. Linear pBR322DNA (Inspiralis) was used at 10.5 nM where indicated. Assays were performed in microtitre plates with a reaction volume of 100 μl. The absorbance at 340 nm was measured continuously in a plate reader(SpectraMAX190, Molecular Devices) and used to evaluate the oxidation of NADH (using an extinction coefficient of 6.22 mM^−1^ cm^−1^), which relates stoichiometrically to the production of ADP.The results were fitted to a Michaelis – Menten plot according to equation 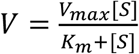 using Origin (Pro), Version 2020 b, (OriginLab Corporation). Novobiocin (50 μM) was used as a control for gyrase-independent ATPase activity, which was found negligible.

### Fluorescence anisotropy measurements

QnrB1 was N-terminally labelled with AlexaFluor 488-carboxylic acid-2,3,5,6-tetrafluorophenyl ester-5 isomer (5-TFP, Thermo), according to a procedure described for YacG protein (32). Briefly, purified QnrB1-His was exchanged to amino labelling buffer (25 mM HEPES-Cl pH 7.5, 10% (v/v) glycerol, 200 mM KCl and 1 mM TCEP, final pH of solution was 7.0). A 15 times molar excess of AlexaFluor 488-5-TFP over QnrB1 was added to the QnrB1 solution. The reaction was incubated in 25°C for 1 h with shaking. Unreacted dye was quenched by adding 1 M L-lysine dissolved in 20 mM Tris-HCl pH 7.5 (final lysine concentration 130 mM) and incubating the reaction at 37°C for 30 minutes. Free dye was separated from labelled protein using Superdex S75 increase 10/300 GL column (Cytiva), equilibrated with QnrB1 storage buffer.

Fluorescence measurements were carried out with a RF-6000 spectrofluorimeter (Shimadzu) fitted with a polarizer. The assays were carried out at specific excitation/emission wavelengths for Alexa-488 of 488/520 nm. For QnrB1 binding to gyrase, 50 nM labelled QnrB1 was mixed in EMSA buffer with appropriate gyrase subunit(s) and incubated for 10 minutes at room temperature in darkness. When 1 mM ADPNP was used, the samples were incubated for additional 1 h at room temperature with addition of 2 mM MgCl_2_. Curves were fitted using Origin (Pro)Version 2020b, (OriginLab Corporation) according to standard one site specific binding equation: 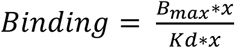 for GyrA and GyrB43. For GyrB, the curve had a different shape and had to be fitted using two sites specific binding equation: 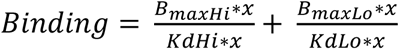

QnrB1/DNA binding competition assays were performed as follows: 90 bp Cy5-labelled oligonucleotide encompassing the strong gyrase binding site from plasmid pBR322 was ordered from Sigma and annealed with an antisense unlabelled oligo by heating to 99°C and gradual cooling. To measure the ability of QnrB1 to compete with DNA, 20 nM 90 bp Cy5-DNA fragment complex was mixed in buffer (50 mM Tris-HCl, pH 7.5, 0.5 mM TCEP, 10 mM MgCl_2_, 30 mM KGlu, 5 % glycerol) with 1 μM gyrase (A_2_B_2_) and increasing concentrations of label-free QnrB1. The assays were carried out at specific excitation/emission wavelengths for Cy5 of 600/666 nm. Optimal concentration of gyrase complex for binding experiments was established beforehands by mixing 20 nM 90 bp Cy5-DNA and increasing concentration of gyrase in above-mentioned buffer. To measure the ability of DNA to compete with QnrB1, 50 nM AlexaFluor488-labelled QnrB1 was mixed with 1 μM gyrase andand increasing concentration of label free linearised pBR322 (Inspiralis). The assays were carried out at specific excitation/emission wavelengths for Alexa-488 of 488/520 nm.

### Pull-down experiments

5 μM purified FLAG-QnrB1 was mixed with 0.65 μM gyrase A_2_B_2_ complex in 25 mM Tris-Cl pH 7.5, 5 mM MgCl_2_, 50 mM KCl and 5% glycerol (buffer also contained ~5 mM arginine hydrochloride and ~5 mM NaCl from QnrB1 storage buffer) in 20 μl volume. For the experiments with ADPNP, the nucleotide (1 mM) was added 30 mins before QnrB1 to induce GyrB dimerisation and the reaction was pre-incubated at RT together with a control reaction. An equal amount of equilibrated FLAG M2 agarose (Sigma) was added to each reaction. After addition of QnrB1, reactions were incubated for a further 60 min at RT with gentle shaking. Resin was washed three times with TGE (20 mM Tris-Cl pH 7.5, 10% glycerol, 100 mM NaCl) and eluted with 50 μg/ml (final concentration) of 3xFLAG peptide (Sigma) in TGE. Load and eluate material were analysed by SDS-PAGE.

### *In vivo* benzoyl-phenylalanine (pBpa)-driven UV crosslinking

*E. coli* GyrA-SPA and GyrB-SPA were co-transformed with pEVOL-pBpa plasmid and pBAD-*QnrB1[x]pBpa* plasmid containing appropriate amber codon substitutions. Cells were grown at 37°C with shaking in LB medium supplemented with antibiotics and 1 mM *p*Bpa. Protein expression was induced by addition of 10 mM arabinose at OD_600_ = 0.6 and culture was allowed to grow for a further 3 hours. After that time cells were centrifuged (5000 g, 10 min), the medium was removed and cells were resuspended in phosphate buffered saline (PBS, pH 7.5). Cell suspensions were poured into plastic Petri dishes and irradiated with UV for 30 min at λ = 365 nm (8 Watt). The temperature of the cell suspension did not exceed 37°C during UV exposure. The samples collected before and after UV-crosslinking were subsequently analysed by SDS-PAGE in 4-20% gradient gels (Bio-Rad TGX). Gels were subsequently transferred onto a PVDF membrane and FLAG epitopes were detected by Sigma M2 antibodies followed by ECL imaging.

### *In vitro* crosslinking experiments

400 nM (final concentration) of relevant gyrase subunits or domains (GyrB/A/B43/B47/B24) and 5 μM (final concentration) of QnrB1*p*Bpa or AlbG*p*Bpa were combined together in EMSA buffer (75 mM KCl, 30 mM Tris-Cl pH 8, 2 mM MgCl_2_, 6% glycerol, 1 mM DTT). After 30 minutes of incubation at room temperature the mixture was irradiated with UV light (λ = 365 nm) in UV-transparent Eppendorf tubes for 30 min. The temperature of the reaction was monitored and did not exceed 37°C. For time-course experiments, aliquots were taken at the indicated time points. For ADPNP containing experiments, 1 mM ADPNP was added 30 mins prior to crosslinking to induce GyrB or GyrB43 dimerization. The reaction products were analysed on SDS-PAGE gel. Competition crosslinking experiments with WT QnrB1 and QnrB1 Y123pBpa were performed as follows: 5 μM (final concentration) of QnrB1*p*Bpa and 400 nM (final concentration) A_2_B_2_ gyrase complex and appropriate amount of WT QnrB1 in EMSA buffer were incubated for 30 minutes in RT. The samples were irradiated with UV light (λ = 365 nm) in UV-transparent Eppendorf tubes for 30 min. The reaction products were analysed on SDS-PAGE gel

### Agar diffusion assay

Agar diffusion assays were performed to investigate the ability of AlbG on its own to sequester/neutralise albicidin. The assay was performed according to the method described elsewhere (41). Briefly an overnight culture of *E. coli* BW25113 was diluted to an OD_600_ = 0.05 with 0.80% LB-agar. AlbG-albicidin reaction mixtures consisted of 40 μM albicidin and appropriate amounts of AlbG (40 μM, 80 μM, 120 μM, 160 μM). Reaction mixtures were incubated for 20 min at room temperature in the darkness. 30 μl aliquots were transferred in triplicates to 2 mm holes made in the inoculated agar plate. Plates were incubated overnight at 37°C and the inhibition zones diameters were measured.

## RESULTS

### Pentapeptide repeat proteins offer specific protection against their cognate toxins *in vivo*

To compare protective activities of different PRPs in identical conditions, we created a set of arabinose-inducible pBAD plasmids, carrying coding sequences for tagless QnrB1, McbG or AlbG. *E. coli* BW25113 (*ara*-) were transformed with one of these plasmids or with an empty pBAD vector, and MICs of CFX, ALB and MccB17 were measured upon induction. Similarly, to the previously published data (24), induction of QnrB1 led to a 16-fold increase of CFX MIC compared to the reference (**Table 1**). Induction of McbG led to a modest 4-fold increase, and no increase in CFX MIC was observed upon AlbG induction. In stark contrast, tests with ALB showed that upon AlbG induction, its MIC increased >128-fold (higher concentrations could not be tested due to the limited solubility of albicidin). Expression of QnrB1 did not increase the albicidin MIC, and expression of McbG resulted in a 4-fold increase of MIC. Finally, the MICs for MccB17 were increased 37-fold for the McbG producing strain, whereas AlbG and QnrB1 induction resulted in 6-fold and 3-fold increase of MIC, respectively. These results clearly indicated that all three PRPs provide specific protection against their cognate toxins.

**Table 1.** MIC data obtained for PRPs and gyrase targeting toxins. **Measured MIC values.** Measured MICs of albicidin (Albi), ciprofloxacin (CFX) and microcin B17 (MccB17) for *E. coli* BW25113 strain transformed with empty vector (pBAD) or plasmids expressing different PRPs.

| Plasmid | Toxin |  |  |  |  |  |
| --- | --- | --- | --- | --- | --- | --- |
|  | Albicidin |  | MccB17 |  | CFX |  |
| | Obtained MIC<br>[μM] | Fold<br>change | Obtained MIC<br>[μM] | Fold<br>change | Obtained MIC<br>[μM] | Fold<br>change |
| pBAD | $0.028 \pm 0.005$ | 1 | $0.58 \pm 0.09$ | 1 | $0.016 \pm 0.004$ | 1 |
| pBAD-QnrB1 | $0.023 \pm 0.008$ | 1 | $1.3 \pm 0.1$ | 2 | $0.25 \pm 0.06$ | 16 |
| pBAD-McbG | $0.13 \pm 0.02$ | 5 | >15 | >26 | $0.12 \pm 0.03$ | 8 |
| pBAD-AlbG | >10 | >341 | $2.6 \pm 0.2$ | 4 | $0.016 \pm 0.004$ | 1 |
| pBAD-AlbG $\Delta_{91-97}$ | $0.062 \pm 0.009$ | 2 | - | - | - | - |

It was previously shown that the deletion of three amino acids (107-109) from QnrB1 loop 2 compromises protection against FQs (21, 24). AlbG contains a similarly located loop (91-97), and the structure of loop deletion Δ_91-97_ AlbG variant has been reported (13), but not tested for protection from albicidin. This prompted us to produce this mutant and evaluate its activity. Expression of Δ_91-97_ *albG* led to a 16-fold lower MIC value for ALB, compared to the WT *albG* gene when tested *in vivo* in our system. (**Table 1**).

### AlbG and QnrB1 rescue *E. coli* gyrase from their cognate toxins by reducing cleavage complex formation

To investigate the biochemical basis of protection offered by PRPs, we purified hexahistidine-tagged QnrB1 and AlbG (we were unable to purify functional McbG in sufficient quantity and purity) and first tested their activities in gyrase supercoiling assays. Similar to previous reports (26), QnrB1 provided limited protection (supercoiling was not completely restored) against 5 μM CFX (**Figure 1C**). Calculated EC50_QnrB1_ (concentration of QnrB1 required to observe half of maximum protective effect) was 0.2 μM. Complete rescue was observed when CFX concentration was lowered to 1 μM. As can be seen from the same figure, high concentrations of QnrB1 (>10 μM) were found to inhibit gyrase activity, and this high concentration inhibitory effect was also observed without CFX present (**Supplementary Figure S1A**). Calculated IC_50 QnrB1_ was 11 μM, >50 times higher than EC50. The abovementioned Δ_107-109_ mutant of QnrB1 (QnrB1 ΔTTR) showed the same level of supercoiling inhibition as WT QnrB1 but was unable to rescue gyrase supercoiling inhibited by CFX. (**Supplementary Figure S1BC**).

When tested against albicidin, AlbG provided a protective effect (supercoiling was restored) over a wide range of concentrations (**Figure 1D**) with EC 50_AlbG_=1.2 μM. In contrast to QnrB1, AlbG did not inhibit *E. coli* gyrase at any concentration tested (up to 50 μM) (**Figure 1E**). Akin to the *in vivo* results, we saw no protection when we swapped the inhibitors and tested protection by QnrB1 against ALB or protection by AlbG against CFX (**Figure 1EF**). Due to the lack of inhibition by AlbG, we wondered whether the mechanism of gyrase protection might be different (e.g. not involving gyrase, such as sequestering of albicidin by AlbG). However, incubation of albicidin with purified AlbG did not show any signs of inactivation of the toxin (**Supplementary Figure S2**).

It was previously shown that QnrB1 reduced the amount of DNA gyrase complexes with cleaved DNA (DNA cleavage) stabilised by CFX (26). We have confirmed this result, showing that in the cleavage assay, the amount of linear DNA in the presence of 5 μM QnrB1 is decreased ~50% across concentrations of ciprofloxacin tested (up to 20 μM) (**Figure 2A, C**). Inhibition of FQ-induced cleavage was also observed when negatively supercoiled DNA was used as a substrate, but the effect was weaker (**Supplementary Figure S3AB**).

**Figure 2.**
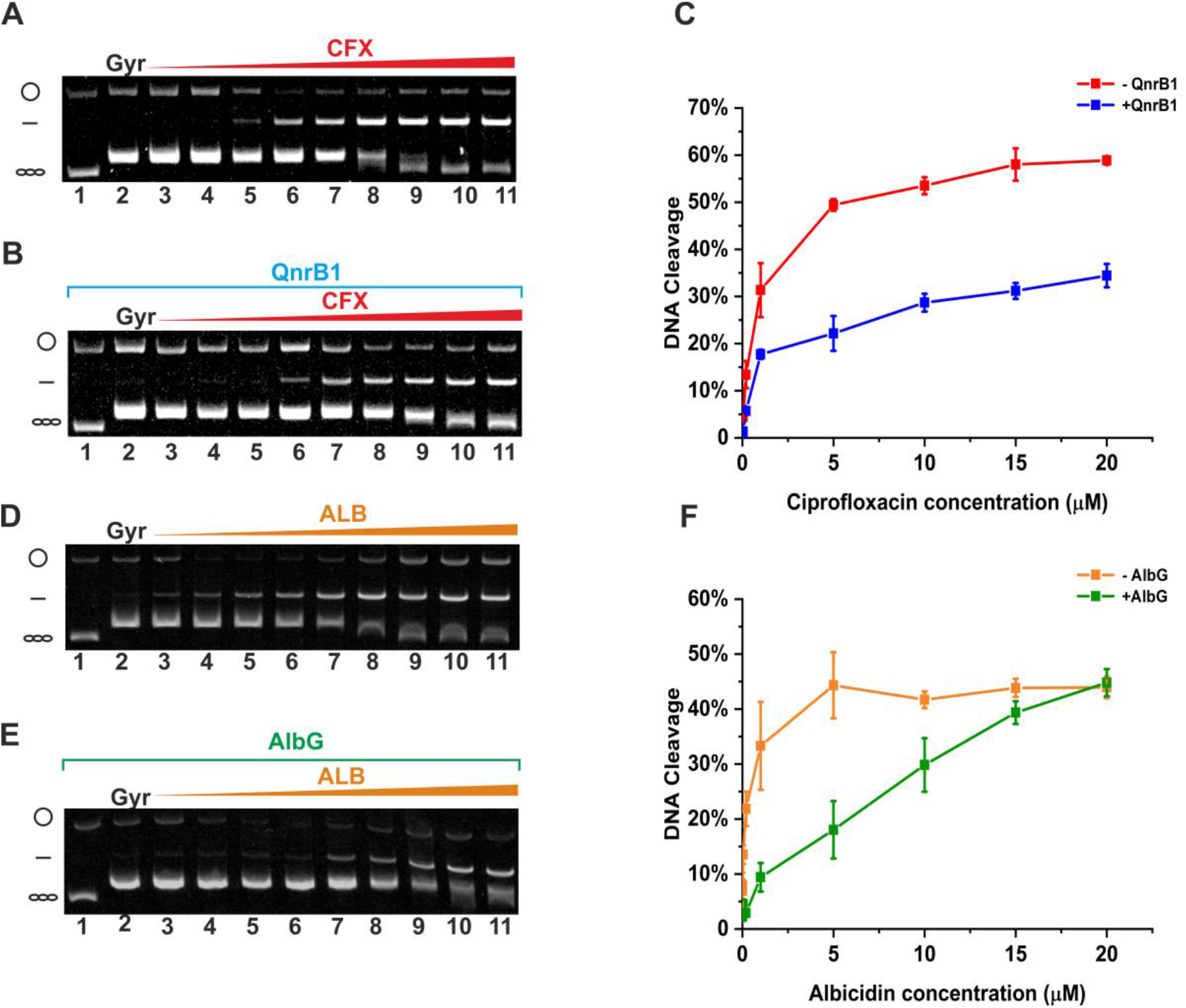
PRPs decrease the amount of cleaved DNA species. Gyrase cleavage assay show that PRP offer protection from cleavage induced by gyrase poisons. (**A**) DNA gyrase (5 U) cleavage reactions with the increasing amount of ciprofloxacin run in the presence of ethidium bromide (EtBr). (**B**) DNA gyrase (5 U) cleavage reactions with the increasing amount of ciprofloxacin in a presence of 5 μM QnrB1, run on EtBr gel. Lane 1, relaxed pBR322. Lane 2, relaxed pBR322 with DNA gyrase. Lanes 3 - 11 contain relaxed pBR322 gyrase and increasing concentrations of ciprofloxacin (0.0016; 0.0008; 0.04; 0.2; 1; 5; 10; 15; 20 μM) (**C**) Cleaved DNA in the absence (*blue*) and presence (*red*) of QnrB1 quantified and plotted. Error bars represent standard deviation (SD) of 3 independent experiments. (**D**) DNA gyrase (5 U) cleavage reactions with the increasing amount of albicidin run on EtBr gel. (**E**) DNA gyrase (5 U) cleavage reactions with the increasing amount of albicidin in a presence of 5 μM AlbG run on EtBr gel. Lane 1, relaxed pBR322. Lane 2, relaxed pBR322 with DNA gyrase. Lanes 3 - 11 contain relaxed pBR322 gyrase and increasing concentrations of albicidin (0.0016; 0,.0008; 0.04; 0.2; 1; 5; 10; 15; 20μM) (**F**) Cleaved DNA in the absence (*red*) and presence (*blue*) of AlbG quantified and plotted. Error bars represent standard deviation (SD) of 3 independent experiments.

AlbG similarly reduced the amount of linear DNA formed in the presence of albicidin (**Figure 2DE**. However, in contrast with QnrB1, here the magnitude of the effect faded with increasing concentration of albicidin, vanishing at [ALB] >15 μM (**Figure 2EF**). As for supercoiling assays, QnrB1 and AlbG showed no cross-protection towards ALB and CFX (not shown).

As GyrB ATPase domains and GyrA CTDs are not required for FQ-stimulated cleavage, we have produced different previously reported truncated gyrase complexes: GyrA_2_59_2_/B_2_ andGyrA_2_/B47_2_ and tested if DNA cleavage by these complexes(using negatively supercoiled DNA as a substrate) can be inhibited by QnrB1. GyrA59_2_/B_2_ complex lacks DNA-wrapping domains (GyrA CTDs) but can relax negatively supercoiled DNA in the presence of ATP, similarly to Topo IV (42). In contrast, GyrA_2_/B47_2_ lacks the GyrB43 ATPase-transducer domain but is still capable of ATP-independent relaxation (43). No protective effect of QnrB1 was seen in cleavage assays with any of these enzymes apart from A59_2_/B_2_ (**Supplementary Figure 3CD**), where the amount of linear DNA trapped by CFX was observed to decrease in the presence of 5 μM QnrB1. Therefore, we assumed that DNA wrapping is not required for the QnrB1 to act, but the ATPase domain is indispensable.

To further investigate the potential role of DNA wrapping, we tested short linear DNA fragments (76, 100, 133, 147, 220 or 300 bp) as a substrate for full-length gyrase cleavage (**Supplementary Figure S4**); again, reduction of cleavage was clearly visible even with the shortest fragment tested, meaning that QnrB1 indeed does not require a DNA node to act.

### QnrB, but not AlbG decreases DNA binding to gyrase

The observed specific protective action of both AlbG and QnrB1 seems inconsistent with the G-segment DNA mimicry model, where PRPs compete with DNA for binding to the enzyme; however, such DNA mimicry could explain the weak inhibitory effect of QnrB1. We carried out EMSA experiments to test whether QnrB1 or AlbG could influence the DNA binding by gyrase. In these assays, QnrB1 decreased the affinity of gyrase to DNA (147 bp fragment, encompassing strong gyrase site from plasmid pBR322) with EC50 = 10.93 ± 0.58 μM, which is very close to the IC50 of 11.25 μM observed in the supercoiling assays (**Supplementary Figure 5AB**). No such effect could be observed for AlbG (tested up to 70 μM, **Supplementary Figure 5C**). In the presence of CFX, which stabilises DNA binding, QnrB1 could not outcompete DNA (**Supplementary Figure S5D**). Additional fluorescence anisotropy experiments using labelled QnrB1 or labelled 90 bp pBR322 DNA fragment also showed competition between linear DNA and QnrB1 (**Supplementary Figure S5EF).** We conclude that G-segment DNA mimicry cannot explain the ability of QnrB1 or AlbG to rescue gyrase supercoiling and inhibit DNA cleavage, manifested at much lower concentrations. However, it can possibly account for the inhibitory effect of QnrB1, observed at high concentrations.

### QnrB1 and AlbG require ATP hydrolysis to rejuvenate poisoned gyrase complexes

G-segment binding and cleavage generally do not require ATP, and thus a hypothetical mechanism of protection, in which PRPs prevent drug binding or promote dissociation of bound drug from the complex, does not require the presence of the nucleotide. However, when we performed cleavage assays without a nucleotide, we found that the CFX-induced cleavage was not reversed by QnrB1 (**Supplementary Figure S6A**). Similar experiments have been done with albicidin, where we tested a range of poison concentrations; again, no protection was observed (**Supplementary Figure S6B**). Therefore, in order for PRPs to act, they require either the dimerization of the ATPase-gate following ATP binding, or ATP hydrolysis and subsequent “resetting” (44) of the enzyme. To choose between these scenarios for QnrB1, we performed time-course experiments to closely monitor CFX-dependent cleavage complex formation in three different conditions: in the presence of ATP, its non-hydrolysable analogue ADPNP or in the absence of nucleotide (**Figure 3AB**). To give QnrB1 an opportunity to bind to the gyrase before N-gate dimerization, nucleotides were added last to the mixture, following preincubation of gyrase and QnrB1. Addition of QnrB1 had no effect on cleavage in case of ADPNP or when nucleotide was omitted; however, in the presence of ATP, a decrease in linear DNA was readily observed, allowing more complete supercoiling in the presence of QnrB1. The negative result with ADPNP suggests that N-gate dimerization by itself is not sufficient for QnrB1 to inhibit cleavage.

**Figure 3.**
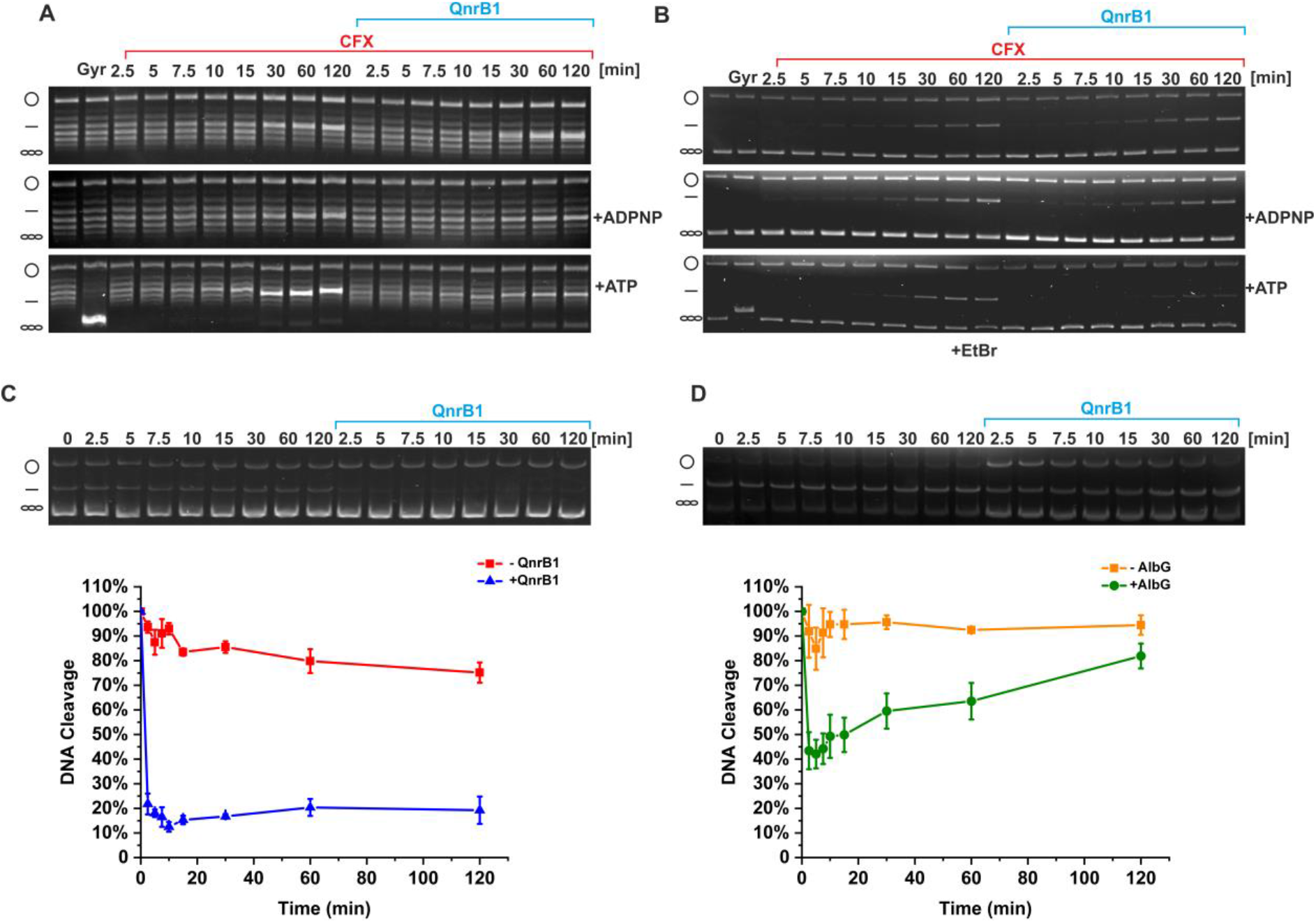
QnrB1 requires ATP hydrolysis to rescue DNA gyrase. (**A**) Time courses of DNA cleavage. The reactions contained 5 μM ciprofloxacin and 5 μM QnrB1 (as indicated) and were run without nucleotide, with ADPNP or ATP. After completion the reactions were run on a gel without EtBr. (**B**) Same reactions run in the presence of EtBr. The amount of cleaved DNA is reduced when 5 μM QnrB1 is present and if ATP was added to the reaction. (**C**) Cleavage complex stability determined in a presence of 5 μM QnrB1. Initial DNA cleavage reactions with 80 U of gyrase and 20 μM ciprofloxacin and were incubated for 10 minutes at 37°C to reach equilibrium and then diluted 20-fold with reaction buffer with or without 5 μM QnrB1. (*Top*) The samples run on EtBr gel. (*Bottom*) Linear DNA was quantified and plotted. Level of DNA cleavage at time 0 was set to 100%. Error bars represent the SD of at least three independent experiments. (**D**) Cleavage complex stability determined in a presence of 5 μM AlbG. Initial DNA cleavage reactions with 80 U of gyrase and 1 μM albicidin were incubated for 10 minutes at 37°C to come to equilibrium and then diluted 20-fold with reaction buffer with or without 5 μM AlbG. (*Top*) The samples ran on the EtBr gel. (*Bottom*) Linear DNA was quantified and plotted. Level of DNA cleavage at time 0 was set to 100%. Error bars represent the SD of at least three independent experiments.

If PRPs merely prevent cleavage complex formation (based on G-segment mimicry or similar mechanism), they must interact with the gyrase before the G-segment is cleaved and covalently bound to the enzyme. Conversely, if PRPs actively promote dissociation of the drugs and DNA re-ligation, they should be able to revert already existing cleavage complexes. We tested if QnrB1 can destabilise pre-formed cleavage complexes, consisting of gyrase, DNA and CFX. **Figure 3C** shows that in the presence of ATP and QnrB1, the pre-formed cleavage complex was quickly dissociated whereas without QnrB1, the complex was stable for at least 2 hours. No dissociation was observed without the nucleotide or with ADPNP (not shown). Thus, QnrB1 is able to interact with gyrase *after* cleavage complex formation to rescue the stalled enzyme at the stage after DNA binding, and this process requires ATP hydrolysis. We have repeated the experiments with the AlbG/albicidin pair and observed the similar effect, but albicidin cleavage complexes appeared more stable in agreement with higher potency of ALB (**Figure 3D**).

### Protective effects of QnrB1 and AlbG do not depend on strand passage

We showed that the inhibition of cleavage complex formation by QnrB1 and AlbG depends on ATP hydrolysis by gyrase. Two possible mechanisms can explain such behaviour: either PRPs require the energy of ATP hydrolysis and associated large-scale conformational changes to achieve removal of bound drug, or ATP-driven strand passage provides PRPs with access to a temporarily exposed drug binding site within the enzyme. The strand passage requirement was suggested previously for MccB17 and gyrase-binding toxin CcdB (40, 41). In these two cases, strand passage, occurring during ATP-independent relaxation of negatively supercoiled DNA, allowed for binding of toxins. We have investigated if ATP-independent relaxation of negatively supercoiled DNA, inhibited by CFX or albicidin, can be rescued by QnrB1 or AlbG. We saw no protection in both cases (**Supplementary Figure S7AB**) but high-concentration inhibitory effects of QnrB1 were still observed, suggesting that QnrB1 is still able to interact with gyrase. Interestingly, AlbG was also found to inhibit gyrase under these conditions.

However, both QnrB1 and AlbG on its own could not inhibit ATP-independent relaxation **(Supplementary Figure S7CD**); instead, QnrB1 slightly promoted it (**Supplementary Figure S7E**). To further test the potential requirement for strand passage, we analysed relaxation activities of A59_2_/B_2_ and A_2_/B47_2_ gyrase complexes mentioned above. Rescue of CFX-poisoned ATP-dependent (A59_2_/B_2_) or ATP-independent (A_2_/B47_2_) relaxation respectively was not observed and the enzyme was strongly inhibited (**Supplementary Figure S8AB**). However, the effects of QnrB1 on its own on ATP-dependent and ATP-independent relaxation were different: while A59/B relaxation was clearly inhibited, relaxation by A/B47_2_ was stimulated akin to the results with the full-length enzyme (**Supplementary Figure S8CD**).

In summary, these observations led us to think that strand passage on its own is not important for QnrB1 activity. Therefore, we hypothesized that QnrB1 interacts with ATPase domains of GyrB, which allosterically leads to the removal of the drug.

### QnrB1 stimulates gyrase ATPase activity

We proceeded to test if QnrB1 and AlbG can stimulate DNA-independent and DNA-stimulated ATPase activity of gyrase. **Figure 4A** shows that QnrB1 indeed increased the ATP hydrolysis rate about 3-fold in the absence of DNA while the DNA-stimulated rate was not affected (K_m − QnrB1_ = 0.46 ± 0.15 μM, K_m + QnrB1_ = 0.96 ± 0.17 μM). Strikingly, when tested with isolated GyrB43 subunit, stimulation became much stronger (**Figure 4BCD**). The ATPase reaction rate of GyrB43 with 5 μM QnrB1 was ~17 times higher than the rate with no QnrB1 present (V_max − QnrB1_ = 0.986 ± 0.075 μmol/min vs V_max + QnrB1_ = 17.1 ± 1.1 μmol/min). Loop deletion mutant QnrB1ΔTTR stimulated ATPase activity equally to the WT variant (**Supplementary Figure S9**).

**Figure 4.**
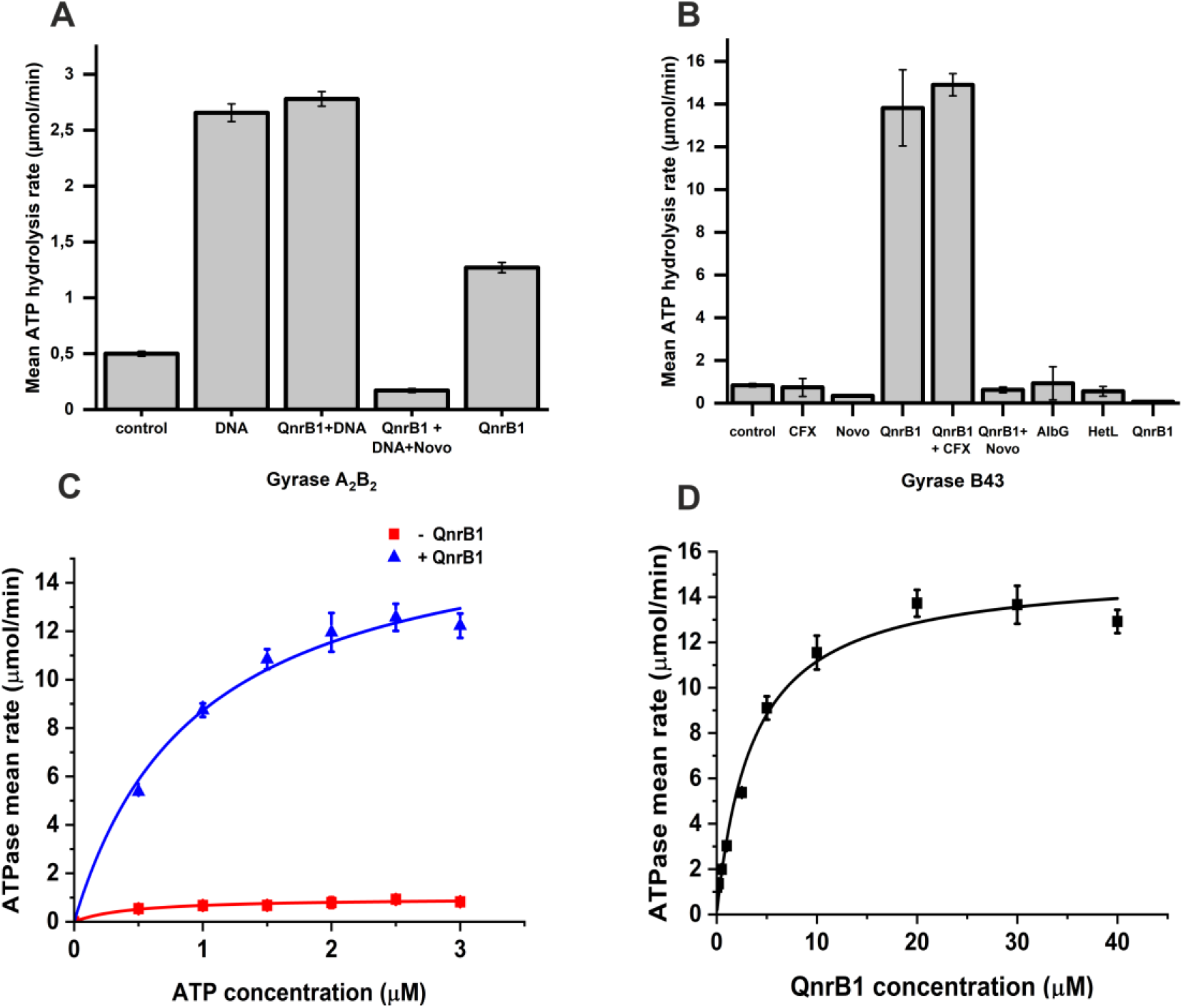
QnrB1 stimulates gyrase ATPase activity. **(A)** ATPase rate data for 50 nM gyrase (A_2_B_2_) complex mixed with: 10 nM DNA; 10 nM DNA and 5 μM QnrB1; 10 nM DNA, 5μM QnrB1 and 50 μM novobiocin; 5 μM QnrB1. **(B)** ATPase rate data for 4 μM GyrB43 mixed with proteins, drugs and DNA as indicated. DNA was used at 10 nM, novobiocin (Novo) at 50 μM, CFX at 5 μM and proteins at 5 μM. **(C)** ATPase rate data and Michaelis-Menten fits for reactions conducted with different concentrations of ATP with (*blue*) or without (*red*) 5 μM QnrB1 for gyrase B43. **(D)** ATPase assays data and Michaelis-Menten fit for reactions with 4 μM GyrB43 conducted with different concentrations of QnrB1 at constant concentration of ATP (1 mM). For all plots, error bars are expressed as the standard deviation of three independent experiments for the rate (μmol/min).

Obtained K_m_ values for ATP in QnrB1-stimulatedGyrB43 reactions lie in the same order of magnitude as values for non-stimulated reactions, suggesting that at the addition of QnrB1 in this concentration does not affect ATP binding (**Figure 4D**). QnrB1-dependent activation of GyrB43 ATPase activity seemed to follow Michaelis-Menten kinetics with K_m_ = 3.61 ± 0.57 μM and V_max_ = 15.19 ± 0.62 μmol/min. (**Figure 4B**). Experiments with full-length GyrB subunit have shown very weak (less than 2-fold) but reproducible activation (not shown). Lastly, we could not observe any stimulation effect for AlbG (**Figure 4B**)

### QnrB1 binds to the GyrB subunit *in vitro*

We sought to determine whether GyrB43 ATPase domain is indeed involved in QnrB1 binding. We measured interactions between gyrase and N-terminally labelled QnrB1 using a fluorescent anisotropy (FA)-based assay, similarly to the work done previously for gyrase regulator YacG (32). As can be seen from **Figure 5A**, [Alexa488]-QnrB1 binds to both GyrB43 and full-length GyrB with estimated K_d GyrB43_ of 1.67 ± 0.31 μM and K_dHi GyrB_ 0.06 ± 0.05 μM respectively. Analysis of GyrB data required fitting to the two-sites binding equaltion (see Materials and methods) which might stem from the complicated nature of interactions between GyrB and QnrB1 dimers. Gyrase A_2_B_2_ complex bound QnrB1 with K_d A2B2_ of 0.08 ± 0.01 μM. Comparing those data with K_d GyrA_: 4 ± 2 μM we concluded that QnrB1 binds specifically within the GyrB subunit. Preincubation of subunits with ADPNP decreased the binding: QnrB1 bound to GyrB with K_dHi_ of 0.16 ± 0.11 μM and to GyrB43 with K_d_ = 18 ± 13 μM. The most significant drop in affinity was observed for A_2_B_2_ complex where K_d_ increased >20-fold to 1.78 ± 0.31 μM (**Supplementary Figure S10**), suggesting that QnrB1 cannot interact with the “restrained” GyrB conformation, stabilised by the nucleotide analog (45).

**Figure 5.**
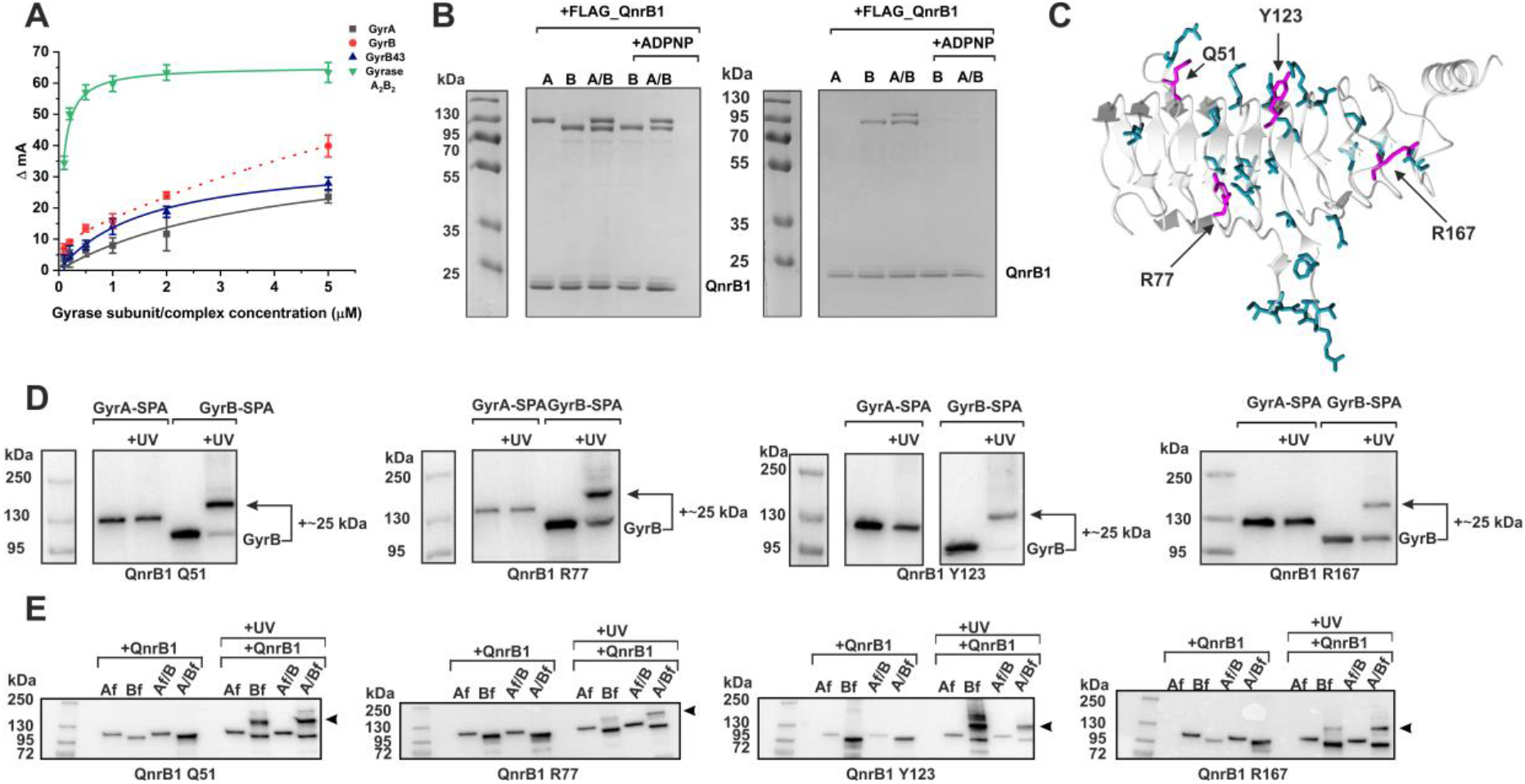
Interactions between GyrB and QnrB1. **(A)** Fluorescence anisotropy assay showing binding of individual gyrase subunits and A_2_B_2_ complex to QnrB1. Solid curves represent one-site binding fits, dotted curve represents postulated two-site binding. Gray square – GyrA; red circle – GyrB; blue triangle – GyrB43; green triangle – gyrase A_2_B_2_ complex. ΔmA indicates the change in anisotropy (in milliunits). Error bars represent the SD of at least three independent experiments. **(B)** *In vitro* pull-down assay with N-terminally FLAG-tagged QnrB1 and purified GyrA, GyrB and A_2_B_2_ complex. *Left* – input, *right* – pull-down (eluates from M2 (α-FLAG) agarose). A Coomassie stained SDS-PAGE gel is shown. **(C)** A QnrB1 monomer is shown in grey cartoon representation with the QnrB1 residues chosen for crosslinking experiments. A cartoon representation of QnrB1 monomer (grey, PDB: 2XTW) is shown with residues chosen for crosslinking experiments displayed as dark cyan cylinders. Residues that produce crosslinks when replaced with *p*Bpa are marked with arrows and displayed as magenta cylinders. **(D)** *In vivo* crosslinking anti-FLAG western blots of QnrB1 Q51, R77, Y123 and R167 *p*Bpa variants to chromosomally encoded GyrA-SPA and GyrB-SPA in *E. coli*. Visible band-shifts correspond to an increase in molecular weight of ~25 kDa, roughly equivalent to QnrB1. **(E)** *In vitro* crosslinking anti-FLAG western blots of QnrB1 Q51, R77, Y123 and R167 *p*Bpa variants with different FLAG-tagged gyrase subunits and mixtures of both tagged and untagged subunits (GyrA/3xFLAG-GyrB or GyrA-FLAG/GyrB) (Af-C-terminally FLAG gyrase A; Bf-N-terminally 3xFLAG gyrase B). Crosslinks are indicated by an arrow.

We used an orthogonal technique (pull-down) to confirm these findings. N-terminally FLAG-tagged QnrB1 was able to bind purified GyrB, both independently and in the context of GyrB_2_A_2_ complex (**Figure 5B**). GyrA on its own didn’t interact with FLAG-QnrB1, but it was retained on the resin in the presence of GyrB, suggesting that gyrase complex is not disrupted by QnrB1 binding. When reactions were pre-incubated with ADPNP, no binding was observed. In the reverse pull-down experiment (**Supplementary Figure S11**), QnrB1 was retained in the presence of 3xFLAG-GyrB, 3xFLAG-GyrB/GyrA complex or GyrB/GyrA-FLAG complex. Once again, preincubation with ADPNP completely abolished binding. We have tried to detect interactions of QnrB1 with smaller parts of GyrB (i.e. ATPase-transducer GyrB43, ATPase GyrB24 and TOPRIM GyrB47) but only full-length GyrB could pull-down QnrB1 (**Supplementary Figure S12**).

### *In vivo* and *in vitro* crosslinking further support QnrB1-GyrB interaction

Attempts to purify QnrB1-gyrase complex by size-exclusion chromatography were unsuccessful, suggesting the transient nature of the complex. We used an orthogonal photo-crosslinkable amino acid benzoyl-phenylalanine (*p*Bpa) to stabilise the native QnrB1-gyrase complex in *E. coli* cells (34). To pinpoint which residues of QnrB1 are likely to be involved, we selected by visual inspection of published crystal structure (PDB:2XTW) 29 surface-exposed QnrB1 residues, which were not part of pentapeptide repeats and did not have obvious structural role, and replaced them with the *p*Bpa (**Supplementary Table S1** and **Figure 5C**). *E. coli* DY330 derivatives, in which chromosomal GyrB or GyrA genes are fused with SPA purification tags (GyrA-SPA and GyrB-SPA) (47, 48) were used as hosts for expression of QnrB1 *p*Bpa variants. Four residues (Q51, R77, Y123, R167) were found to produce crosslinks, all of them with GyrB (**Figure 5D**, **Supplementary Table S1** and **Supplementary figures 18-20**). All crosslinked residues were found on one face of QnrB1 (Face 2), suggesting a defined interaction interface (**Figure 5C**).

In order to confirm these results, crosslinking experiments were repeated *in vitro* with purified Q51*p*Bpa, R77*p*Bpa, Y123*p*Bpa and R167*p*Bpa QnrB1 variants and FLAG-tagged *E. coli* gyrase subunits described above. Q51*p*Bpa and Y123*p*Bpa were crosslinking strongly to GyrB while R77*p*Bpa and R167*p*Bpa gave weaker crosslinks, but also to GyrB (**Figure 5E**). Molecular weight of the main crosslinked band corresponded to the attachment of a single QnrB1 molecule to GyrB. Additional shifted species were observed as weaker bands, corresponding to the attachment of two QnrB1 molecules. Interestingly, while the double crosslink band was very strong for GyrB-Y123*p*Bpa, it completely disappeared in case of the GyrA/B complex, suggesting that only the single crosslink band is biologically relevant.

Additional experiments with Q51*p*Bpa and Y123*p*Bpa QnrB1 variants and purified GyrB subdomains showed that QnrB1 was almost quantitatively crosslinked to GyrB and to the GyrB43 domain (**Supplementary Figure S13**) but did not interact with a shorter GyrB24 domain or TOPRIM (GyrB47) (though a very weak band was detected for TOPRIM-Y123*p*Bpa).

To further prove the specificity of observed crosslinks, we have carried out competition experiments, where increasing amounts of WT QnrB1 were added to the crosslinking reactions with GyrB and QnrB1 Y123*p*Bpa or GyrB43 and QnrB1 Y123*p*Bpa (**Supplementary Figure S14**). In both cases, WT QnrB1 easily replaced QnrB1 Y123*p*Bpa, with 10x excess of unlabelled protein completely preventing crosslinking.

To support our hypothesis that PRP-gyrase interactions are conserved, we have produced an AlbG mutant (AlbG N109*p*Bpa) which is structurally equivalent to QnrB1 Y123*p*Bpa and carried out *in vivo* and *in vitro* crosslinking experiments (**Supplementary Figure S15 and Supplementary Figure S16**). Similarly, to QnrB1, AlbG D109*p*Bpa was shown to specifically crosslink to GyrB subunit in the context of GyrA/B complex, suggesting that at least some interactions between PRPs and gyrase are conserved.

Following the results of FA and pull-down experiments where ADPNP was found to block the GyrB-QnrB1 interaction, we have tested the effects of nucleotide analog-induced dimerisation on the observed crosslinks. We performed *in vitro* crosslinking experiments with QnrB1 Y123*p*Bpa and GyrB43 or full-length GyrB pre-incubated with ADPNP to induce subunit dimerization. ADPNP did not influence the crosslink efficiency of GyrB43 (**Supplementary Figure S17B**), but largely prevented crosslinking of full-length GyrB subunit with QnrB1 (**Supplementary Figure S17A**). Importantly, the QnrB1 ΔTTR mutant (Y123pBpa Δ_107-109_) was equally well crosslinked to GyrB (**Supplementary Figure S17C**).

## DISCUSSION

### Specificity of PRP interactions with toxins

The G-segment DNA mimicry model originally proposed for MfpA postulated that this protein acts by reducing DNA binding to gyrase, which limits the formation of toxic cleavage complexes. The proposed reduction in DNA binding should reduce susceptibility to all agents that stabilise cleavage complexes involving the G-segment-gyrase interface. However, Qnr does not protect against proteinaceous gyrase poison CcdB (45) nor against natural product simocyclinone D8, which binds to the GyrA subunit in the ‘saddle’ region and prevents G-segment DNA binding. Moreover, QnrB1 was found to act synergistically with simocyclinone (47). In our work we show that the activity of three different PRPs, QnrB1, AlbG and McbG show a high level of specificity towards three different gyrase poisons namely CFX, albicidin and MccB17. QnrB1 provided maximum protection against CFX and had almost no effect on the action of albicidin or microcin B17. Similarly, AlbG was extremely specific to albicidin. These results were recapitulated *in vitro* with purified gyrase. Given the overall structural similarity between different topoisomerase-interacting PRPs, a universal higher level mechanism of action would be expected, however this should allow for specific interactions depending on the particular gyrase poisons.

### Effects of QnrB1 and AlbG on supercoiling and relaxation reactions of DNA gyrase

Topoisomerase-interacting PRPs were proposed to be gyrase regulators with different activities. *Mycobacterium tuberculosis* MfpA, *Klebsiella pneumoniae* Qnr and *Enterococcus faecalis* EfsQnr were all reported to both protect *E. coli* gyrase from quinolone inhibition and at the same time inhibit gyrase supercoiling activity. In this study we tried to carefully disentangle “inhibition” and “rescue” modes of action of QnrB1 using *E. coli* gyrase as a model. In supercoiling reactions, QnrB1 required >10000-fold excess over the enzyme for efficient inhibition. The same level of inhibition was observed with a loop mutant protein, devoid of any protective activity. In contrast, AlbG from *Xanthomonas albilineans* did not produce any visible inhibition of supercoiling even at >10000-fold excess over gyrase.

DNA gyrase is capable of ATP-independent relaxation of negatively supercoiled DNA, where strand passage is believed to proceed in the reverse direction (‘bottom to top’). Strikingly, this reaction was not inhibited by QnrB1 at any concentration (in fact, we consistently observed slight stimulation of relaxation activity at the highest concentrations of QnrB1 - see **Supplementary Figure S7**). Similarly, ATP-independent relaxation of negatively supercoiled DNA by GyrA/GyrB47 complex was not inhibited but rather stimulated by high doses of QnrB1 (**Supplementary Figure S8**). (see below)

In the case of ATP-dependent, active relaxation by the GyrA59/GyrB complex strand passage is supposed to happen ‘top to bottom’ from N-gate to the C-gate (42). This reaction was strongly inhibited by QnrB1. Therefore, QnrB1 is only able to inhibit gyrase activity which requires normal “top to bottom” strand passage, coupled with ATP hydrolysis.

Lastly, we did not observe any inhibitory activity for AlbG, which comes from a Gram-negative bacterium *X. albilineans*. *Xanthomonas* gyrase has a significantly lower homology with the *E. coli* enzyme compared to the *Klebsiella* gyrase (59% versus 92% for GyrA and 61% versus 95% for GyrB). We hypothesize that AlbG has much stronger binding to its cognate *Xanthomonas* enzyme, as required for the host organism protection against a very potent toxin. It is worth noting that *Xanthomonas* gyrase was reported to have significantly different enzymatic properties and multiple antibiotic resistance (48).

### Inhibition of DNA binding by QnrB1/AlbG

The main assumption of the “G-segment DNA mimicry” hypothesis is that PRPs bind to gyrase and reduce the amount of bound DNA. The only direct evidence for this was reduced DNA binding in filter assays with a Qnr variant (26). We have directly tested the ability of QnrB1 and AlbG to prevent DNA binding in fluorescence anisotropy (FA) and gel retardation (EMSA) experiments. EMSAs with 147 bp linear pBR322 fragments do show a decrease of DNA binding when QnrB1 was added to the reaction mix, however, the effect was visible only at concentrations of QnrB1 ~10-fold higher than required for protection (IC50=~11 μM). Other protein inhibitors, directly occluding the DNA binding site on gyrase, such as YacG are reported to have apparent Ki as low as 35 nM (32). Moreover, in the presence of CFX which stabilises DNA binding, QnrB1 was unable to outcompete DNA. Similar results were obtained in FA experiments. This behaviour is expected given that QnrB1 does not inhibit the normal function of gyrase unless at very high concentrations.

Finally, no effect on DNA binding was found for AlbG, despite the latter protein’s clear ability to relieve albicidin inhibition. We suggest that in line with the results above showing that AlbG has poor affinity for E. *coli* gyrase, which is nevertheless sufficient to rescue it from albicidin inhibition.

### Rescue of poisoned gyrase complexes by QnrB1 and AlbG

We have shown that a significantly reduced excess of QnrB1 over gyrase (100-fold) is required to effectively relieve FQ inhibition, compared with the >10000-fold excess required to inhibit gyrase. A slightly higher working concentration was required for AlbG (200-fold excess over gyrase), while no inhibition was observed for this protein as discussed above. Being the first to characterise AlbG activity *in vitro* against its cognate gyrase poison, we have additionally shown that it does not act by sequestering albicidin, analogous to the recently described AlbA (41). We did not see any signs of inactivation of the toxin after incubation with AlbG, confirming that direct interaction with the gyrase is the most likely mode of action for AlbG. Both QnrB1 and AlbG decreased CFX- or albicidin-induced cleavage complex formation respectively. The concentration of CFX did not affect the ability of QnrB1 to inhibit cleavage complex formation, whilst AlbG failed to efficiently protect gyrase at [ALB] higher than 15 μM, which is not surprising given the differences between *X. albilineans* and *E. coli* gyrase discussed above. The cleavage inhibition activity strictly required ATP hydrolysis by gyrase, ruling out any models where PRPs are expected to simply bind to the enzyme and block the gyrase poison binding site. Moreover, we have shown that both QnrB1 and AlbG are able to actively disrupt pre-formed gyrase cleavage complexes in the presence of ATP (but not the non-hydrolysable analog ADPNP). In our view, these findings are hard to reconcile with the G-segment DNA mimicry model. These results are better aligned with the hypotheses suggesting specific recognition of topoisomerase poisons by PRPs (13) or T-segment DNA mimicry theory (see below).

In the absence of ATP, QnrB1 still binds to gyrase-FQ complexes, as manifested by inhibition of DNA relaxation reactions in the presence of FQs. The same is true for AlbG which was found to inhibit ATP-independent relaxation, but only in the presence of albicidin. Therefore, ATP hydrolysis is required for protection, but not for binding of PRPs to the gyrase.

### Interactions of QnrB1 and AlbG with DNA gyrase

Both QnrB1 and MfpA were reported to bind *E. coli* gyrase in native gel retardation (26) and surface plasmon resonance (20) experiments, respectively. Models were proposed for PRPs (20, 23) to bind to the positively charged GyrA dimer. In the more recent pull-down experiment (49), GST-tagged QnrB1 protein was shown to bind *E. coli* GyrB much more strongly than GyrA. Likewise, in two-hybrid system experiments (50) the QnrB1 interaction signal for GyrB was 7- to 11-fold higher than the signal for GyrA. The same work has shown that GyrA, but not GyrB interaction depends on the presence of the QnrB1 loops and is affected by sublethal doses of FQs.

Our study presents overwhelming evidence that the main binding partner of QnrB1 is GyrB. First, FA experiments have shown that labelled QnrB1 in the absence of DNA binds strongly to isolated GyrB, GyrB43 and especially to the GyrB/A complex, but not to GyrA. Similarly, only GyrB or gyrase A_2_B_2_ complex, but not GyrA alone was pulled down by FLAG-tagged QnrB1 and in the reverse experiment, only GyrB or GyrB-containing complexes were able to pull down QnrB1. Lastly, to map the binding interface, we tested 30 positions across the surface of QnrB1 by site-specific UV-crosslinking first *in vivo* and then *in vitro*. Robust crosslinking with GyrB and GyrB43 was observed with 4 residues, revealing a specific surface on QnrB1 (Face 2) most likely interacting with GyrB. We have confirmed the specificity of this interaction in two ways: 1) in the presence of both GyrA and GyrB, only crosslinking to GyrB was observed; 2) unlabelled QnrB1 was able to outcompete QnrB1-BpA, preventing crosslinking.

In all three types of experiments we performed (FA, pull-downs, crosslinking) pre-incubation of GyrB or GyrA/B with ADPNP prevented QnrB1 binding. Reduction of binding was also observed in FA assays with GyrB43/ADPNP. We propose that the QnrB1 dimer makes extensive contacts along a significant portion of GyrB, interacting with both GyrB43 and TOPRIM in a way that at least a part of QnrB1 enters the cavity formed by TOPRIM domains. GyrB dimerisation leads to steric clashes and precludes binding. However, crosslinking of QnrB1 to GyrB43 was not affected by the presence of nucleotide analogue, meaning that at least a part of the interacting surface is still available in the GyrB43 dimer. We propose that the “open” conformation (51, 52) of gyrase, stabilised by FQs, likely allows for QnrB1 binding. We also cannot exclude that binding of QnrB1 requires GyrB to adopt a new conformation, not seen previously in crystal structures. Any structure-based model of PRP binding should account for these observations.

Pull-down experiments did not provide evidence that AlbG can bind strongly to *E. coli* gyrase. This aligns well with the lack of inhibition of gyrase-catalysed reactions and DNA binding by AlbG compared with QnrB1. Nevertheless, the AlbG D109*p*Bpa variant designed based on QnrB1 Y123*p*Bpa also crosslinked to GyrB, suggesting that, globally, the interactions observed might be universal for all PRPs. We predict that stronger binding would be observed if AlbG was used in tandem with its host *Xanthomonas* gyrase and we plan to test this experimentally in future work.

### Stimulation of ATPase activity by QnrB1

The association between QnrB1 and GyrB is additionally supported by the observed stimulation of ATP hydrolysis. This stimulation is particularly strong in the case of GyrB43 (17-fold). In contrast, only modest (3-fold) stimulation was observed for the full-length gyrase, compared with 7-fold stimulation by DNA. The difference in the magnitude of the QnrB1 effect might be attributed to the much higher baseline ATPase activity, exhibited by the full-length GyrB subunit, compared to GyrB43 (we used 4 μM of GyrB43 versus 50 nM of full-length subunits to achieve comparable reaction rates). QnrB1 cannot stimulate ATPase activity of the full-length gyrase to the same extent as DNA; at the same time, it stimulates ATP hydrolysis of GyrB43. Therefore, QnrB1 is unlikely to bind to the GyrB in exactly the same way as DNA. One possibility is that QnrB1 induces dimerisation or stabilises the GyrB43 dimer and thus promotes ATP hydrolysis.

No stimulation of ATPase activity was observed for AlbG. Again, this is in line with the absence of binding to GyrB, lack of gyrase inhibition, and inability to compete with DNA, suggesting that all these effects are connected to the stronger binding of QnrB1 to the *E.coli* gyrase.

### Role of loops in the activity of QnrB1 and AlbG

A 12-amino acid loop, protruding from the rod-like QnrB1 scaffold, was shown to be the main determinant of protection against FQs (21, 24), with a deletion of three amino acids (107-109, TTR) completely abolishing protection (24). A similar structural element has been found in AlbG (13). Interestingly, in two-hybrid system experiments, loop deletions did not perturb the interaction with GyrB. This corresponds well with our findings that QnrB1 ΔTTR was found to bind GyrB equally strongly as WT protein in pull-down and crosslinking experiments and activated ATPase activity to the same extent. Moreover, QnrB1 ΔTTR inhibited gyrase supercoiling activity to the same extent as WT protein, despite having no visible protective activity. In this study we also for the first time analysed the role of the AlbG loop and found that similarly to QnrB1, its deletion abolished protection against albicidin *in vivo*.

We assume that the loop, as was suggested before (50) is important for the precise positioning of the PRP, required to remove the bound drug, but does not by itself constitute the main binding interface (see below).

### T-segment mimicry model as a possible mechanism of action of topoisomerase acting PRPs

Given the significant structural similarities between PRPs, we expect that the overall mechanism of topoisomerase protection must be universal but should account for the observed toxin-specificity. Moreover, it should account for the two clearly distinct modalities of PRP action (general interaction with the enzyme and drug-specific protection). This is highlighted by the example of the QnrB1 ΔTTR mutant, which is still able to bind gyrase, stimulate ATP hydrolysis and cause inhibition of supercoiling at high concentration, but is completely devoid of protective activity (21, 24). In contrast, AlbG, retaining its loop, does not bind *Ec* gyrase well, but retains the capacity to protect the enzyme against gyrase poison.

The suggestion that PRPs are DNA mimics is attractive, but the original “G-segment” DNA mimicry model is not supported by our and others’ data (21). We have previously hypothesized (31) that topoisomerase targeting PRPs function via a T-segment mimicry mechanism, in which they are captured and translocated through the enzyme in a manner analogous to the T-segment. In such a model PRP would presumably enter the inner cavity delineated by the GyrB43 subunits, in a manner analogous to the short DNA duplex found in a crystal structure of *S. pneumoniae* Topo IV (PDB:5J5Q). In this position PRP would not be expected to make any contacts with the TOPRIM domain, and its binding should not be affected by the GyrB dimerization. The results of FA, pull-down and crosslinking experiments performed in this work, contradict this notion: while ADPNP “locking” does not completely prevent GyrB43-QnrB1 interaction (as the crosslink is still observed), the Kd is markedly increased and all interactions with full-length GyrB are completely abolished. Therefore, QnrB1 (and by extension other PRPs), while mimicking a T-segment in terms of capture and, potentially passage through the enzyme, differ from a T-segment in the details of their interactions with GyrB, namely they are expected to bind in a way to interact with both GyrB43 and TOPRIM domains of GyrB. Interactions with TOPRIM are supported also by the results of A_2_/B47_2_ relaxation assays, where the effects of QnrB1 presence can be consistently seen both with and without FQs.

We propose an alternative model for QnrB1 action which incorporates elements of the” T-segment DNA mimicry model” (**Figure 6**) and suggest that the same model applies to all topoisomerase-interacting PRPs. In this model, to bind to the gyrase in physiological conditions (i.e. without gyrase poison present and with DNA wrapped around the enzyme) QnrB1 must outcompete T-segments, which requires very high excess of QnrB1 over gyrase and is unlikely to happen. However, sufficient QnrB1 interferes with normal top to bottom strand passage and inhibits supercoiling and ATP-dependent relaxation, as observed in our experiments with QnrB1 (**Figure 6**, “Gyrase inhibition”). The same interference helps to promote ATP-independent relaxation, which is thought to occur in the reverse direction, i.e. DNA entering from the C-gate. Inhibition of top to bottom strand passage thus helps to shift the thermodynamic relaxation equilibrium towards more relaxed DNA species. Binding of QnrB1 also destabilises the wrapped DNA complex, reducing DNA binding observed in EMSAs and FA assays. The binding of QnrB1 to gyrase does not require loops, but likely requires amino acids on Face 2.

**Figure 6.**
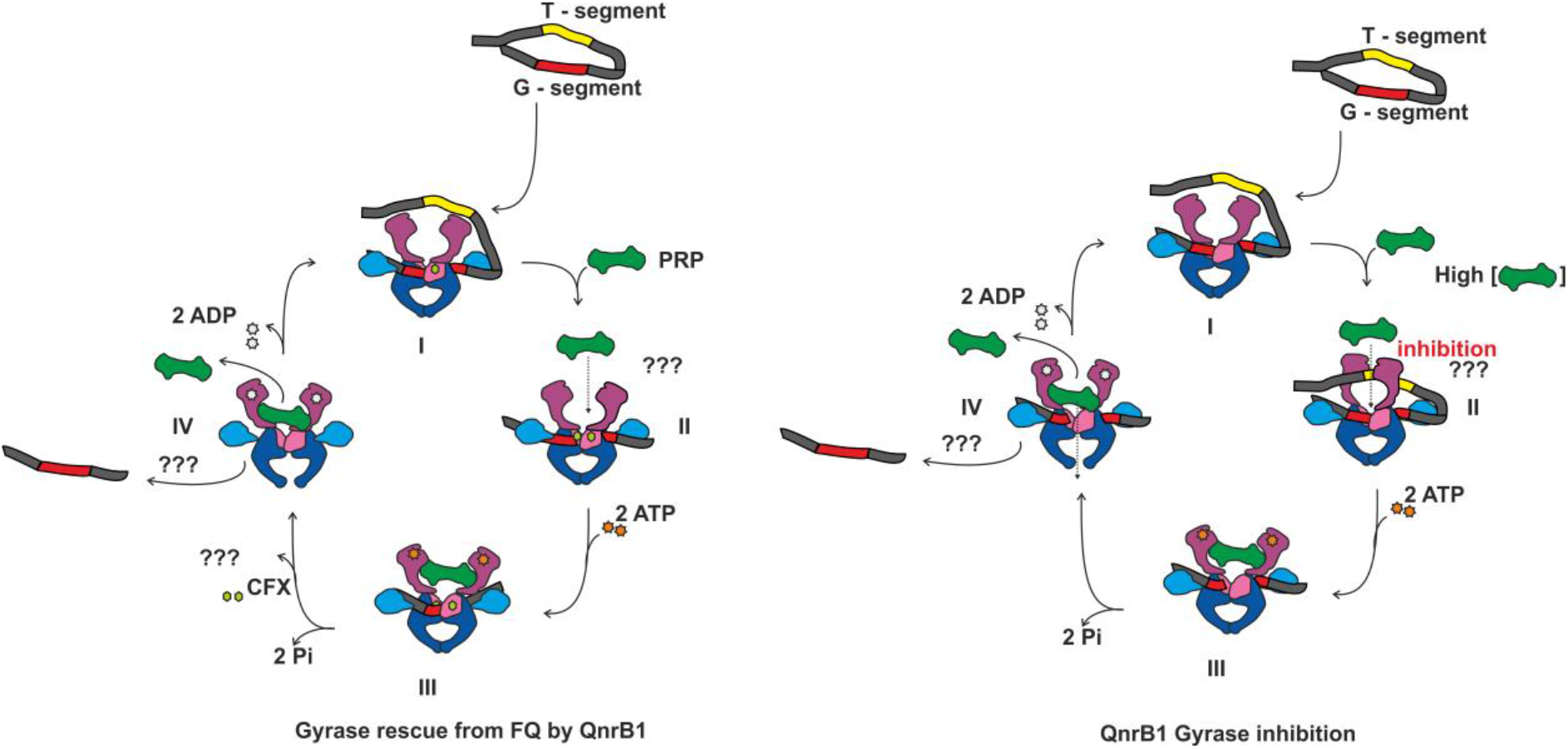
Potential mechanism of action of PRPs (updated T-segment mimicry model). *Left-Gyrase rescue from FQ by QnrB1:* **I**. DNA gyrase cleavage complex formation; **II**. QnrB1 initial binding to the ATP-operated clamp (ATPase and transducer domains) in GyrB. **III, IV**. After ATP binding and hydrolysis specific (loop-mediated) QnrB1 interaction results in fluoroquinolone removal and subsequent release of PRP. This can be accompanied by the DNA release. *QnrB1 gyrase inhibition*: **I**. DNA gyrase with bound G segment; **II**. At high concentrations of PRP it can compete with the T-segment and bind to the ATP-operated clamp, preventing T-segment binding. **III, IV**. After ATP binding and hydrolysis, QnrB1 interaction with gyrase results in the release of PRP, which can be accompanied by the release of DNA.

When the DNA is not present, the QnrB1 does not have to compete with the T-segment and can bind gyrase. Cleavage complex stabilisation by the gyrase poison should similarly allow QnrB1 to interact with the enzyme, presumably by removing competition with the T-segment. The protective activity of PRPs strictly requires ATP hydrolysis by gyrase, moreover, QnrB1 binding actively promotes ATP hydrolysis. If we consider that the protective mechanism involves replacement of bound gyrase poison by the QnrB1 molecule then to efficiently replace bound toxin, the QnrB1-stabilised conformation must have lower energy than the toxin-stabilised one. Energy input is then required for a subsequent QnrB1 release, otherwise the enzyme would be constantly inhibited by the QnrB1 itself. Therefore, the requirement for the energy input in the form of ATP is logical. We suggest that QnrB1/AlbG act as “spokes in the wheel” which, upon ATP hydrolysis, are “pushed” through the enzyme and physically dislodge bound drugs or toxins (**Figure 6**, “Gyrase rescue from FQ”). This dislodgement is poison-specific and requires specific amino acids in loops of QnrB1 or AlbG. Analogous mechanisms of transient interaction with the enzyme are observed for ribosome protective factors such as TetO and TetM which hydrolyse GTP in order to release themselves from the ribosome (53)

Further clarification of this model requires knowledge of fine molecular details of interactions between PRPs and their targets. High-resolution structures of QnrB1 “caught in the act” will be difficult to obtain, as they require full complex assembly and likely ATP hydrolysis. No structures of the full enzyme with T-segment captured at the moment of strand passage have been reported to date. However, the recent development of a cryo-EM platform for structural studies of the *E. coli* gyrase complex (54) has the potential to address such challenges. Such structural information may also allow to design novel gyrase inhibitors based on PRP peptides, novel antibacterials which avoids PRP-driven resistance or small molecules which will block the PRP-gyrase interactions, to keep the potency of existing drugs such as FQs.

## Supporting information

Supplementary Data - includes Supplementary Figures S1-S20 and Supplementary Tables S1-S3

## ACKNOWLEDGEMENTS

Some of the ideas in this work were based on initial observations of Dr. Mikhail Metelev working under the supervision of Dr. Svetlana Dubiley and Prof. Konstantin Severinov. We want to thank Prof. Anthony Maxwell, Prof. David Lawson and Dr. Lipeng Feng for critical reading and comments on the manuscript and for sharing results prior to publication.

