## Supplementary Data - includes Supplementary Figures S1-S20 and Supplementary Tables S1-S3 for "Pentapeptide repeat proteins QnrB1 and AlbG require ATP hydrolysis to rejuvenate poisoned gyrase complexes"

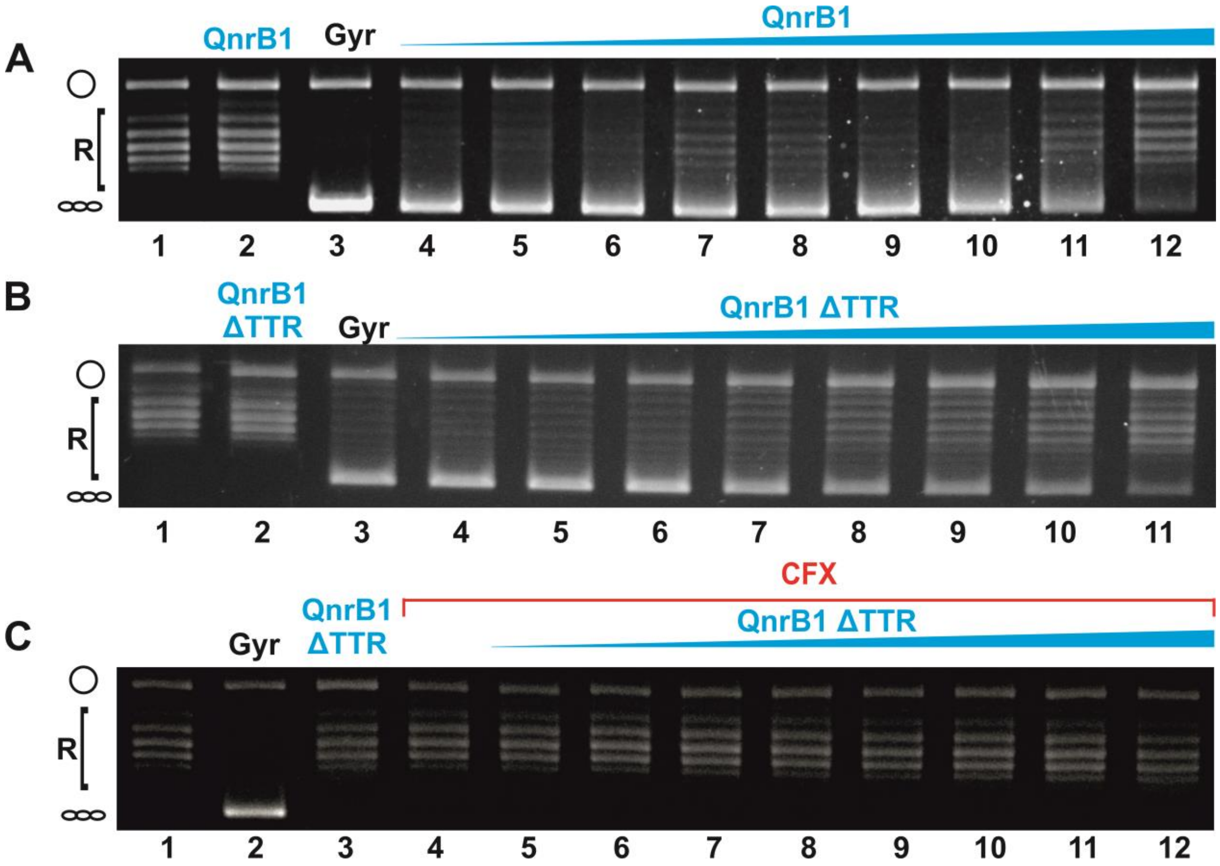


**Supplementary Figure S1. Inhibition of gyrase supercoiling reaction by the increasing amounts of QnrB1.** (**A**) Plasmid supercoiling assay showing the inhibitory effect of high concentrations of QnrB1. Lane 1: relaxed pBR322, lane 2: no nuclease activity was observed upon addition of 50 μM QnrB1, Lane 3: gyrase and relaxed pBR322 lanes 4-12: effect of increasing concentration of QnrB1 on gyrase supercoiling activity (0.0016; 0.008; 0.04; 0.2; 1; 5; 10; 20; 40 μM). (**B**) Plasmid supercoiling assay showing inhibitory effect of high concentrations of QnrB1 ΔTTR. Lane 1: relaxed pBR322, lane 2: nuclease control (pBR322+50 μM QnrB1 ΔTTR); lane 3: gyrase and relaxed pBR322; lanes 4-11: effect of increasing concentration of QnrB1 on gyrase supercoiling activity (0.0016; 0.008; 0.04; 0.2; 1; 5; 10; 25; 50 μM). (**C**) Plasmid supercoiling assay showing lack of protection by QnrB1 ΔTTR. Lane 1: relaxed pBR322; lane 2: nuclease control (pBR322+50 μM QnrB1 ΔTTR); lane 3: gyrase, 1 U; lane 4: gyrase+5 μM CFX; lanes 5-12: gyrase+5 μM CFX +increasing concentration of QnrB1 ΔTTR (0.0016; 0.008; 0.04; 0.2; 1; 5; 10; 25; 50 μM)

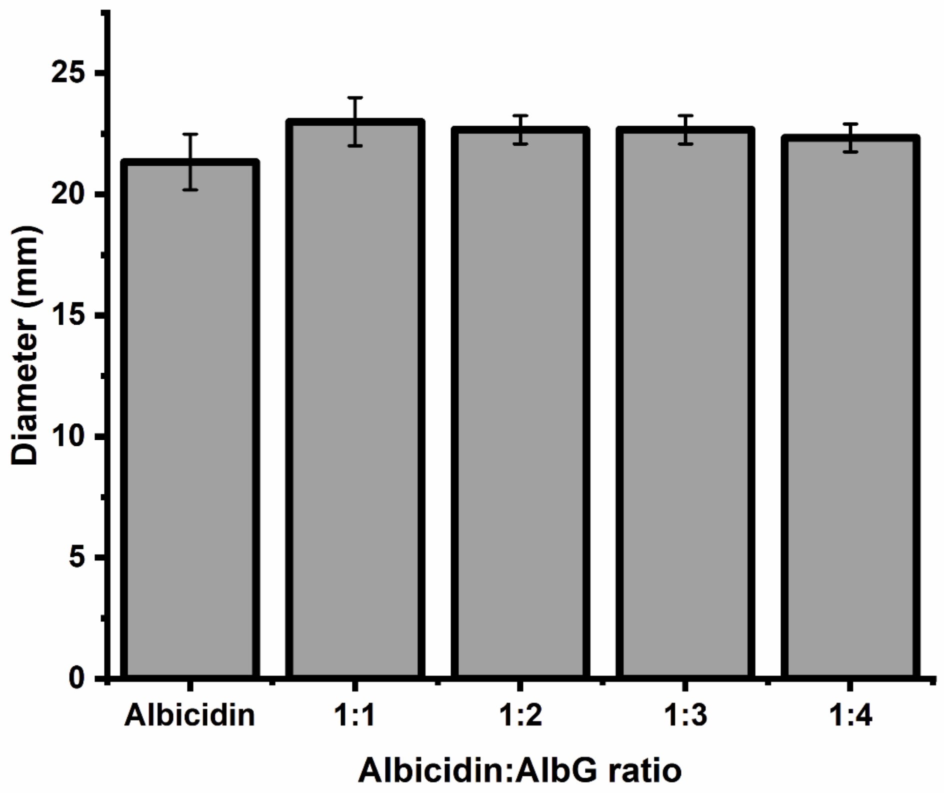


**Supplementary Figure S2. Growth inhibition zones diameter measured for albicidin incubated with different amounts of AlbG.** 40 μM albicidin was mixed with AlbG and transferred to plates inoculated with a tester strain. Error bars are expressed as the standard deviation of three independent experiments.

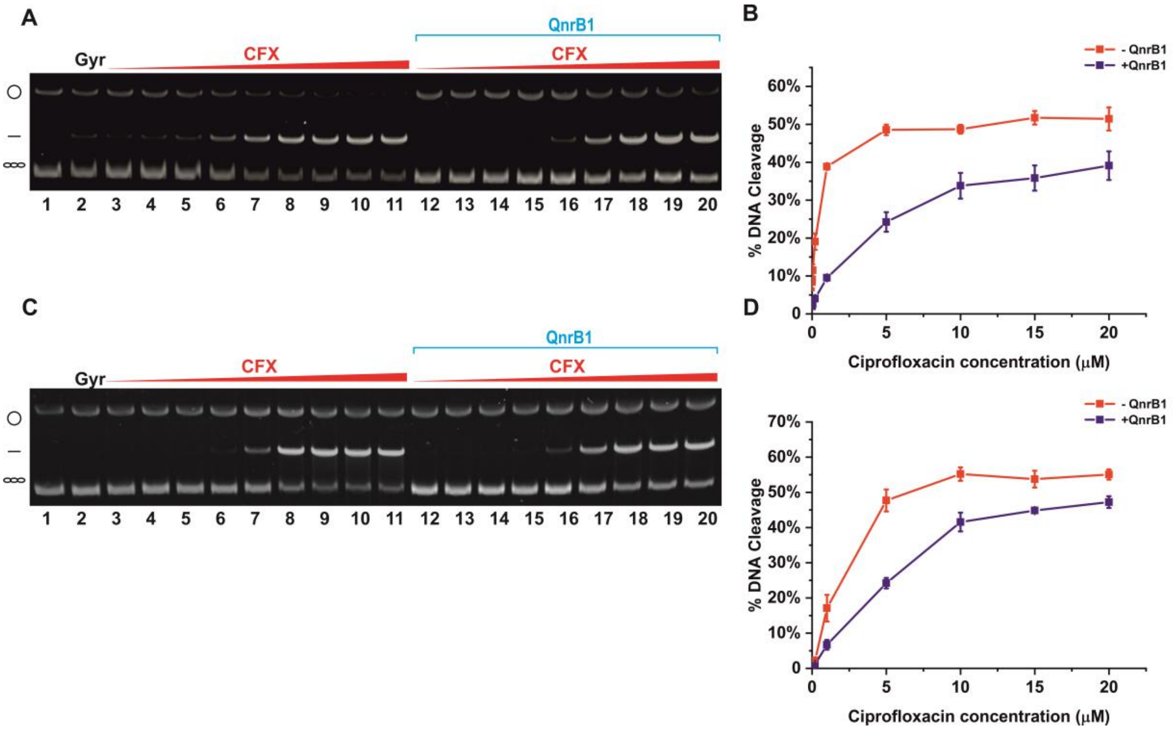


**Supplementary Figure S3. Inhibition of ciprofloxacin-induced gyrase cleavage activity by QnrB1.** (**A**) DNA cleavage assay with gyrase $A_2B_2$ complex. Lane 1: negatively supercoiled pBR322; lane 2: negatively supercoiled pBR322 and 5 U (20 nM) $A_2B_2$; lanes 3 – 11: effect of increasing concentration of ciprofloxacin on DNA cleavage (0.0016; 0.008; 0.04; 0.2; 1; 5; 10; 15; 20 µM). Lanes 12 – 20: effect of increasing concentration of ciprofloxacin on DNA cleavage (same concentrations) in the presence of 5 µM QnrB1. (**B**) Graph showing dependence of DNA cleavage (percent) from ciprofloxacin concentration for $A_2B_2$ with (blue triangles) and without (red dots) QnrB1. Error bars represent standard deviation (SD) of 3 independent experiments. (**C**) DNA cleavage assay with gyrase $A59_2B_2$ complex. Lane 1: negatively supercoiled pBR322; lane 2: negatively supercoiled pBR322 and 20 nM gyrase $A59_2B_2$ complex; lanes 3 – 11: effect of increasing concentration of ciprofloxacin on DNA cleavage (0.0016; 0.008; 0.04; 0.2; 1; 5; 10; 15; 20 µM). Lanes 12 – 20: effect of increasing concentration of ciprofloxacin on DNA cleavage (0.0016; 0.008; 0.04; 0.2; 1; 5; 10; 15; 20 µM) in presence of 5 µM QnrB1. (**D**) Graph showing dependence of DNA cleavage (percent) from ciprofloxacin concentration for $A59_2B_2$ with (blue triangles) and without (red dots) QnrB1. Error bars represent standard deviation (SD) of 3 independent experiments.

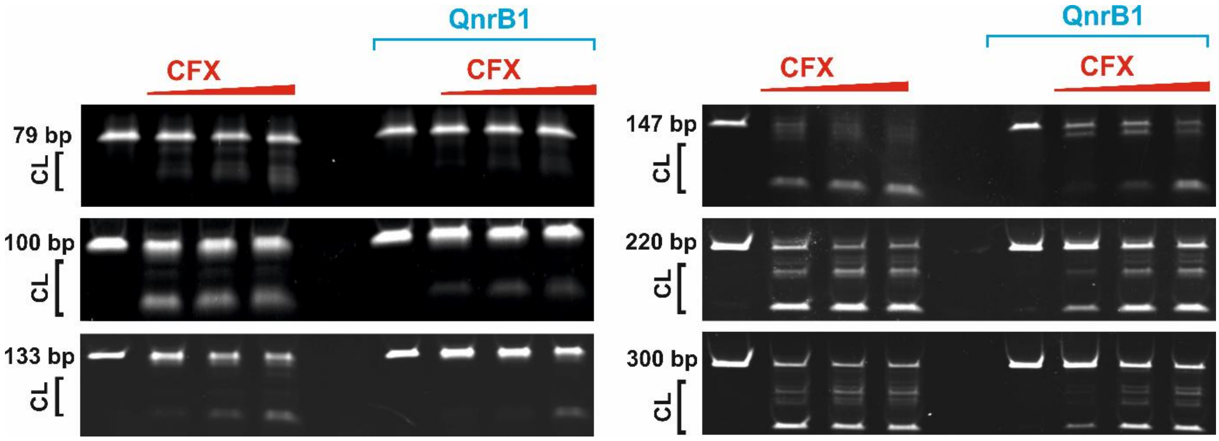


**Supplementary Figure S4**. **Cleavage protection by QnrB1 of DNA fragments of different length at different ciprofloxacin concentrations**. Length of fragments (in base pairs) is indicated at the left. CL – cleaved DNA. 50 nM gyrase complex is present in each reaction. QnrB1 (5 µM) and CFX (0.1, 1, 10 µM) were added as indicated.

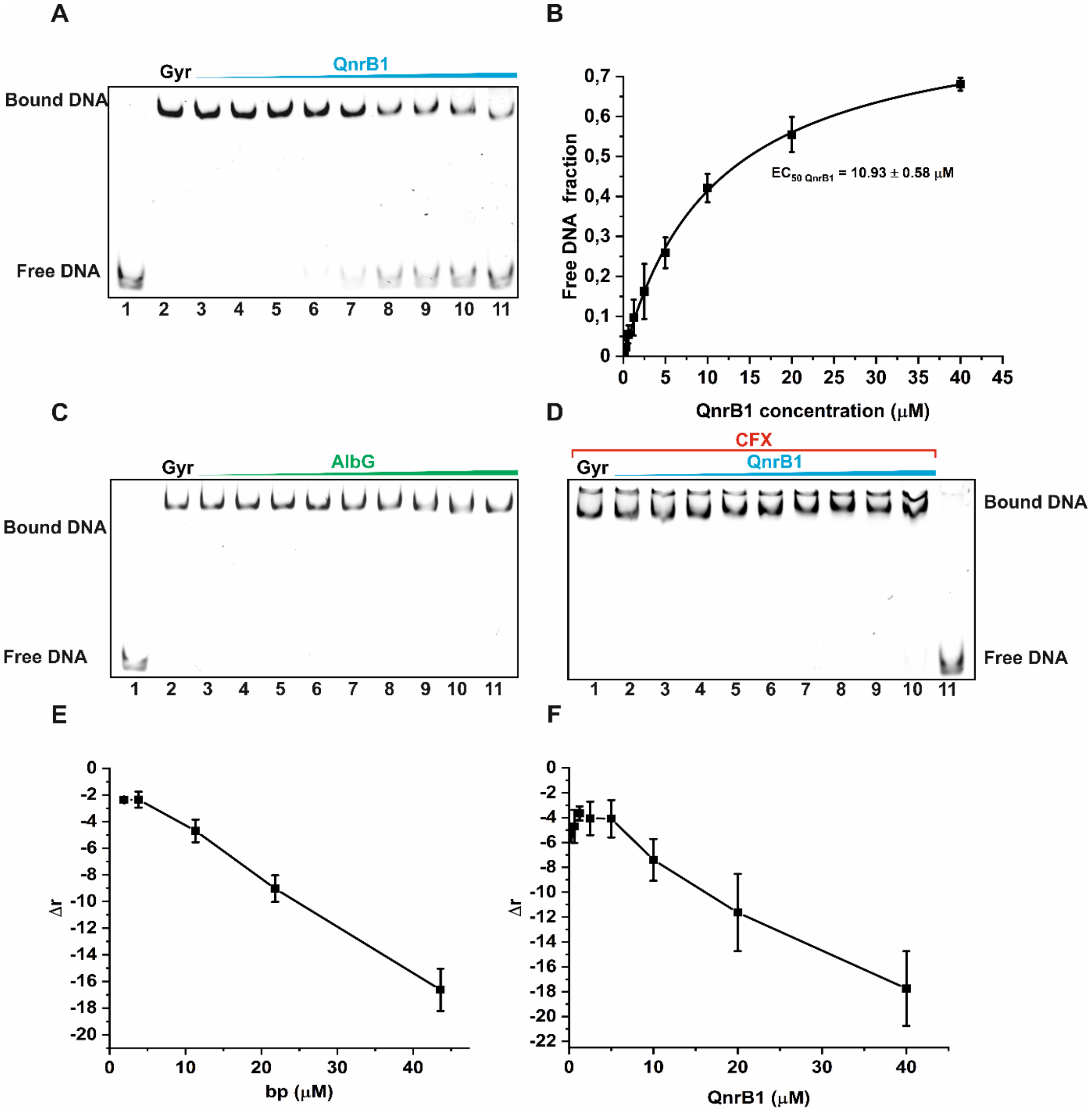


**Supplementary Figure S5**. **Effects of QnrB1 and AlbG on DNA binding by gyrase.** (**A**) EMSA showing the effect of increasing concentration of QnrB1 on DNA binding. Shown is SYBR Gold-stained 6% TBM gel. Lane 1: free 147 bp DNA; lane 2: gyrase (200 nM) added to DNA; lanes 3-11: effect of increasing concentrations (0.2; 0.3; 0.6; 1.3; 2.5; 5; 10; 20; 40 μM) of QnrB1 on DNA binding by gyrase. (**B**) Graph showing dependence of free DNA fraction on concentration of QnrB1. $EC_{50}$ value for QnrB1 and DNA competition is calculated from the fitted curve. Error bars are expressed as the standard deviation of three independent experiments. (**C**) Effect of increasing concentration of AlbG on DNA binding. Lane 1: free DNA; lane 2: gyrase added to DNA; lanes 3-11: effect of increasing concentrations of AlbG(0.1; 0.2; 0.5; 1; 2; 4; 12; 35; 70 μM) on DNA binding by DNA gyrase. (**D**) Effect of increasing concentration of QnrB1 on DNA binding in a presence of 5 μM CFX. Lane 1: gyrase, DNA and 5 μM CFX; lane 2-10: effect of increasing concentration (0.2; 0.3; 0.6; 1.3; 2.5; 5; 10; 20; 40 μM) of QnrB1 on DNA binding by DNA gyrase; lane 11: free DNA. (**E**) Fluorescence anisotropy measurement of QnrB1 displacement by increasing concentration of linear DNA (linear pBR322). 50 nM Alexa-488-QnrB1 and 1 μM DNA gyrase complex were used in reaction. Concentrations of linear DNA calculated as concentration of base pairs (bp) were as follows: (2; 4; 11; 22; 44 μM) (**F**) Fluorescence anisotropy measurement of DNA displacement by increasing concentration of QnrB1. 20 nM 90 bp DNA-Cy5 and 1 μM DNA gyrase complex were used in reaction. Concentrations of QnrB1 were as follows: (0.3125; 0.625; 1.25; 2.5; 5; 10; 20; 40 μM).

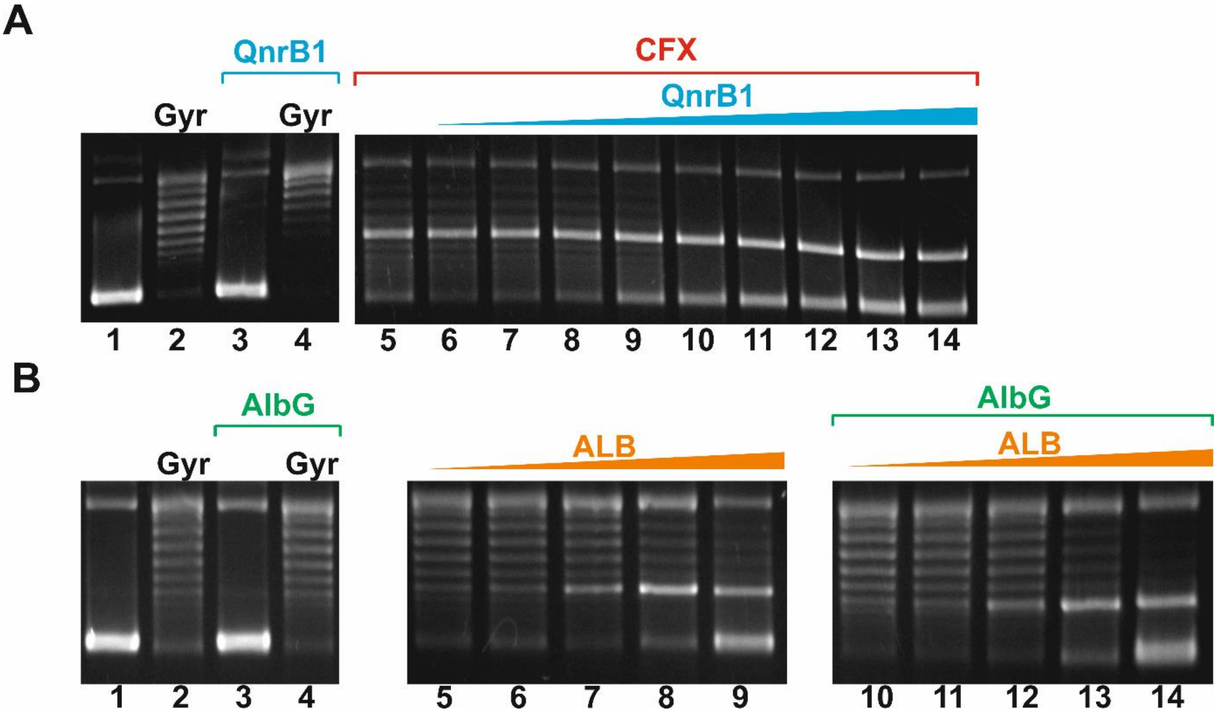


**Figure S6**. **Nucleotide dependence of QnrB1 and AlbG protective activity.** (**A**) Gyrase cleavage assay on negatively supercoiled DNA without nucleotide present in the mixture (in relaxation conditions). (*left*) lane 1: negatively supercoiled pBR322; lane 2: relaxation by gyrase (20 nM); lane 3: nuclease control (pBR322 + purified QnrB1 50 μM); lane 4: pBR322+gyrase+50 μM QnrB1; (*right*) lanes 6-14: DNA cleavage stimulated by CFX (5 μM) and the effect of increasing concentrations of QnrB1 (0.008; 0.04; 0.2; 1; 5; 10; 20; 25; 50 μM). (**B**) Different concentrations of ALB (indicated) were tested together with 10 μM AlbG in a plasmid relaxation assay. Lane 1: negatively supercoiled pBR322; lane 2: relaxation by gyrase (20 nM); lane 3: lack of detectable nuclease activity in the purified AlbG (50 μM); lane 4: 50 μM AlbG + gyrase; (*right*) lanes 5-9: DNA cleavage by gyrase with increasing concentration of ALB (lanes 10-14) and the effect of addition of increasing concentrations of AlbG (0.016; 0.16; 1.6; 16; 160 μM)

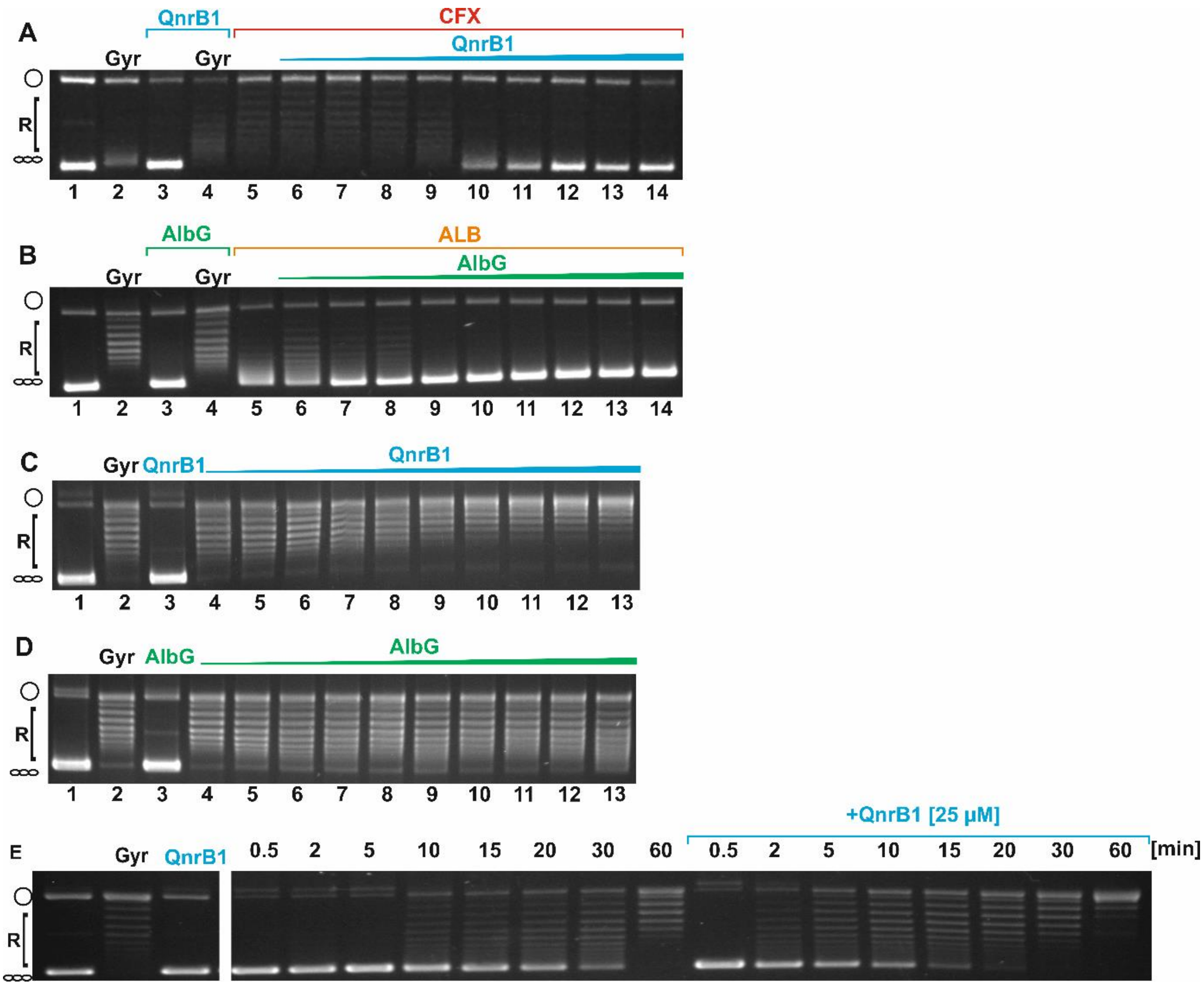


**Supplementary Figure S7**. **Effects of QnrB1 and AlbG on ATP-independent relaxation of negatively supercoiled DNA. (A)** Relaxation of negatively supercoiled DNA in a presence of ciprofloxacin and increasing QnrB1 concentrations. Lane 1: negatively supercoiled pBR322; lane 2: relaxation by 20 nM gyrase; lane 3: lack of detectable nuclease activity in the purified QnrB1 (50 µM QnrB1); lane 4: relaxation in the presence of 50 µM QnrB1; lanes 5-14: DNA relaxation in the presence of CFX (5 µM) and the effect of increasing concentrations of QnrB1 (0.008; 0.04; 0.2; 1; 5; 10; 20; 25; 50 µM). (**B**) Relaxation of negatively supercoiled DNA in a presence of albicidin and increasing AlbG concentrations. Lane 1: negatively supercoiled pBR322; lane 2: relaxation by gyrase; lane 3: lack of detectable nuclease activity in the purified AlbG (50 µM AlbG); lane 4: relaxation in the presence of 50 µM AlbG; lanes 5-14: DNA relaxation in the presence of ALB (5 µM) and the effect of increasing concentrations of AlbG (0.008; 0.04; 0.2; 1; 5; 10; 20; 25; 50 µM). (**C**) Relaxation of negatively supercoiled DNA in presence of increasing QnrB1 concentrations. Lane 1: negatively supercoiled pBR322; lane 2: relaxation by gyrase; lane 3: relaxation in the presence of 50 µM QnrB1; lanes 4-13: relaxation in presence of increasing concentrations of QnrB1 (0.0016, 0.008; 0.04; 0.2; 1; 5; 10; 20; 25;

50 μM). (**D**) Relaxation of negatively supercoiled DNA in presence of increasing AlbG concentrations. Lane 1: negatively supercoiled pBR322; lane 2: relaxation by gyrase; lane 3: relaxation in the presence of 50 μM AlbG; lanes 4-13: relaxation in the presence of increasing concentrations of AlbG (0.0016; 0.008; 0.04; 0.2; 1; 5; 10; 20; 25; 50 μM). (**E)** Time course of DNA relaxation in presence of QnrB1. Gyr: DNA + gyrase after 60 minutes. QnrB1: nuclease control (50 μM QnrB1+DNA). Time points are indicated above the lanes.

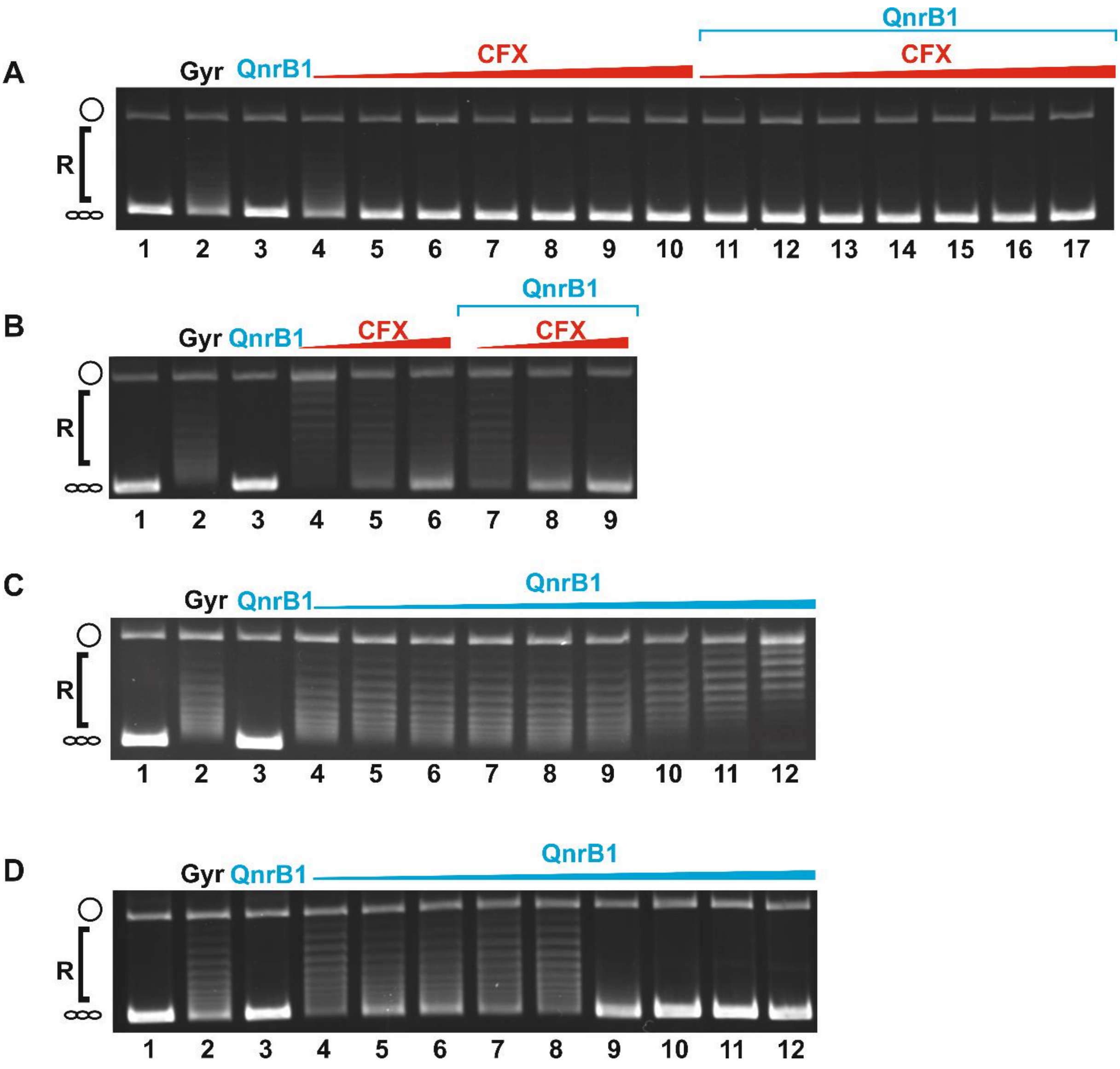


**Supplementary Figure S8. Effects of QnrB1 on relaxation by truncated gyrase enzymes: ATP-dependent (by $A59_2/B_2$) and ATP-independent (by $A_2/B47_2$).** (**A**) $A59_2/B_2$ ATP-dependent relaxation assay in presence of increasing concentration of ciprofloxacin and 5 µM QnrB1. Lane 1: negatively supercoiled pBR322; lane 2: relaxation by $A59_2/B_2$; lane 3: relaxation in the presence of 50 µM QnrB1; lanes 4-10: DNA relaxation in the presence of increasing concentrations of CFX (0.2; 1; 5; 10; 20; 25; 50 µM); lanes 11-17: DNA relaxation in the presence of increasing concentrations of CFX (as before) and 5 µM QnrB1. (**B**) $A_2/B47_2$ ATP-independent relaxation assay in the presence of increasing concentrations of ciprofloxacin and 5 µM QnrB1. Lane 1: negatively supercoiled pBR32; lane 2: relaxation by $A_2/B47_2$; lane 3: relaxation in the presence of 50 µM QnrB1; lanes 4-6: DNA relaxation in the presence of increasing concentrations of CFX (20; 35; 50 µM) ; lanes 7-9: DNA relaxation in the presence of increasing concentrations of CFX (as before) and 5 µM QnrB1 (**C**) $A_2/B47_2$ ATP-independent relaxation assay in the presence of increasing concentrations of QnrB1. Lane 1: negatively supercoiled pBR322; lane 2: relaxation by $A_2/B47_2$; lane 3: relaxation in the presence of 40 μM QnrB1; lanes 4-12: DNA relaxation in the presence of increasing concentrations of QnrB1 (0.0016; 0.008; 0.04; 0.2; 1; 5; 10; 20; 40 μM) (**D**) $A59_2/B_2$ ATP-dependent relaxation assay in the presence of increasing concentrations of QnrB1. Lane 1: negatively supercoiled pBR322; lane 2: relaxation by $A59_2/B_2$; lane 3: relaxation in the presence of 40 μM QnrB1; lanes 4-12: DNA relaxation in the presence of increasing concentrations of QnrB1 (0.0016; 0.008; 0.04; 0.2; 1; 5; 10; 20; 40 μM)

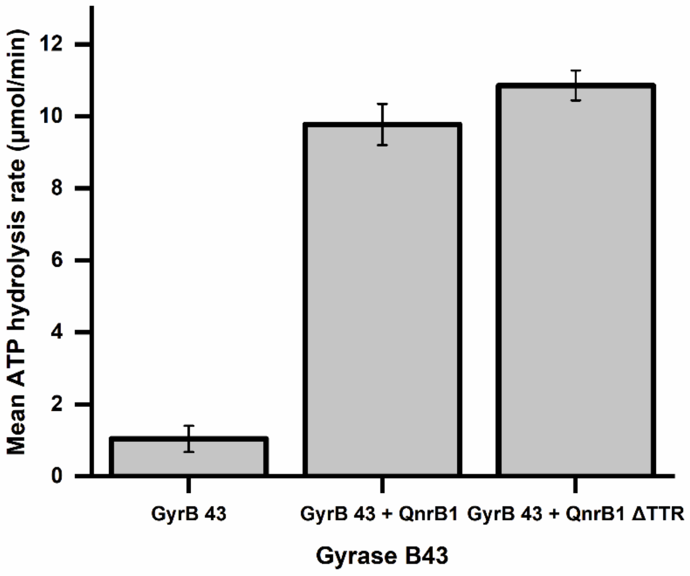


**Supplementary Figure S9. ATPase rate data for QnrB1 ΔTTR.** ATPase rate data for 4µM GyrB43 mixed with 5 µM QnrB1 and 5 µM QnrB1 ΔTTR. Error bars are expressed as the standard deviation of three independent experiments for the rate (µmol/min).

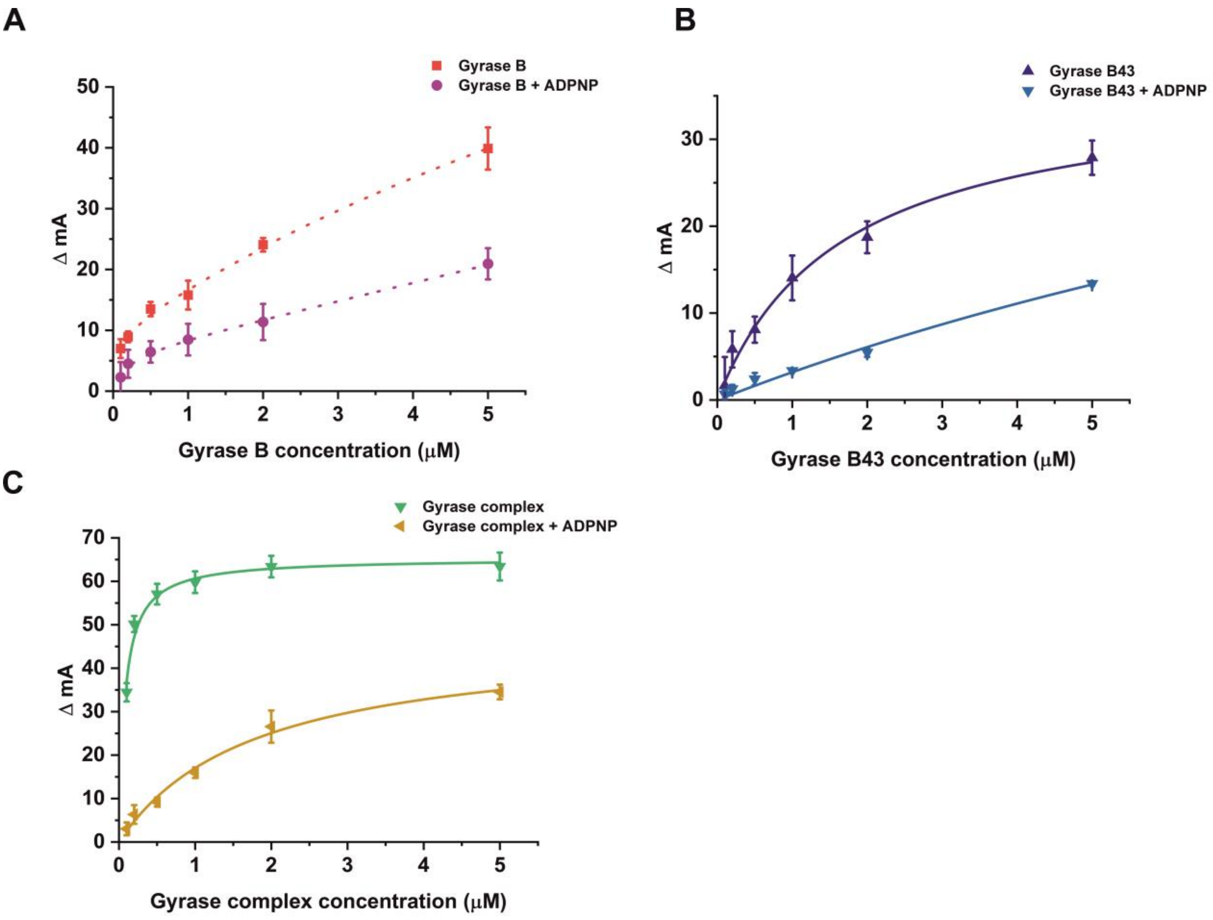


**Supplementary Figure S10. Interaction of gyrase subunits with QnrB1 measured by fluorescence anisotropy.** 50 nM Alexa-488-labelled QnrB1 was used in all experiments. (**A**) GyrB, dotted curves represent fits to a two-sites binding equation ($K_{dHi}$ = 0.06 ± 0.05 μM; $K_{dHi\,(+ADPNP)}$ = 0.16 ± 0.11 μM); (**B**) GyrB43, solid curves represent single-site binding fits ($K_d$ = 1.67 ± 0.31 μM, $K_{d\,(+ADPNP)}$ was 18 ± 13 μM); (**C)** Gyrase $A_2B_2$ complex, solid curves represent one-site binding fits ($K_d$ = 0.08 ± 0.01 μM, $K_{d\,(+ADPNP)}$ = 1.78 ± 0.31 μM). ΔmA indicates the change in anisotropy (in milli - units). Error bars are expressed as the standard deviation of three independent experiments.

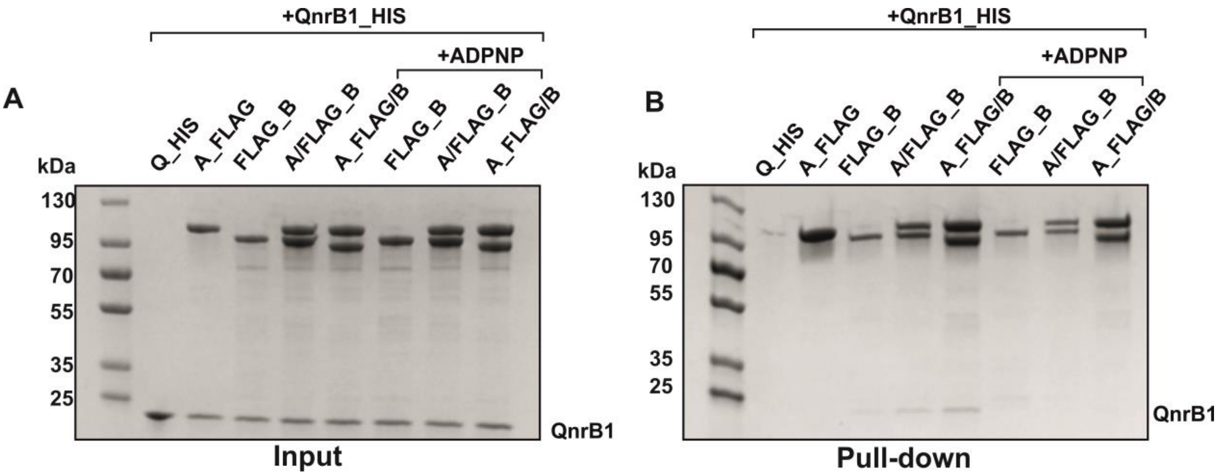


**Supplementary Figure S11. SDS-PAGE gels showing reverse pull down of QnrB1 and different gyrase subunits.** 5 µM QnrB1 and 0.65 µM of gyrase subunit/complex were used in each reaction. Samples were pre-incubated with 1 mM ADPNP where indicated.

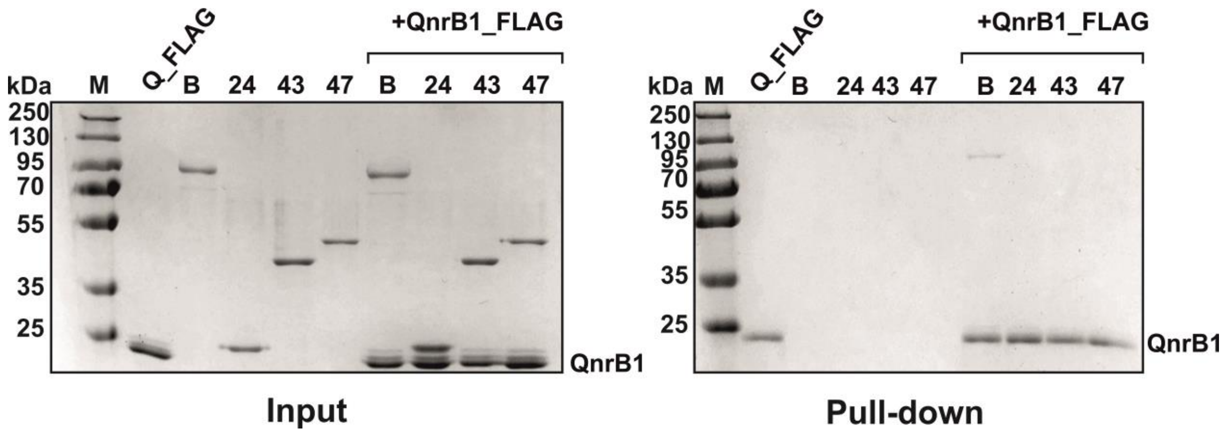


**Supplementary Figure S12. S12 SDS-PAGE gel showing pull-down with FLAG_QnrB1 and GyrB sub-domains.** 5 μM QnrB1 and 0.65 μM of gyrase protein were used in each reaction.

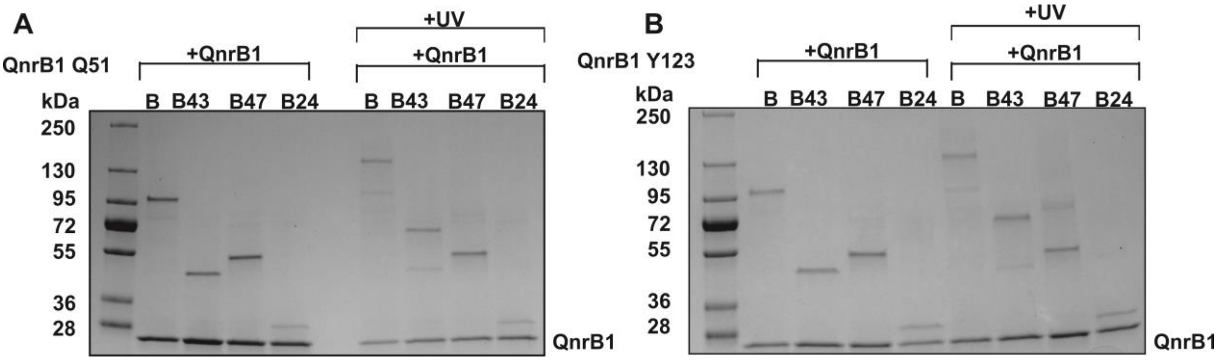


**Supplementary Figure S13. SDS-PAGE gels showing UV – induced crosslinking of QnrB1 Q51 *p*Bpa and QnrB1 Y123 *p*Bpa with different gyrase subdomains.** 5 μM of QnrB1 and 0.4 μM of gyrase subunit as indicated was used in the reaction. (**A**) crosslink with QnrB1 Q51 *p*Bpa; (**B**) crosslink with QnrB1 Y123 *p*Bpa.

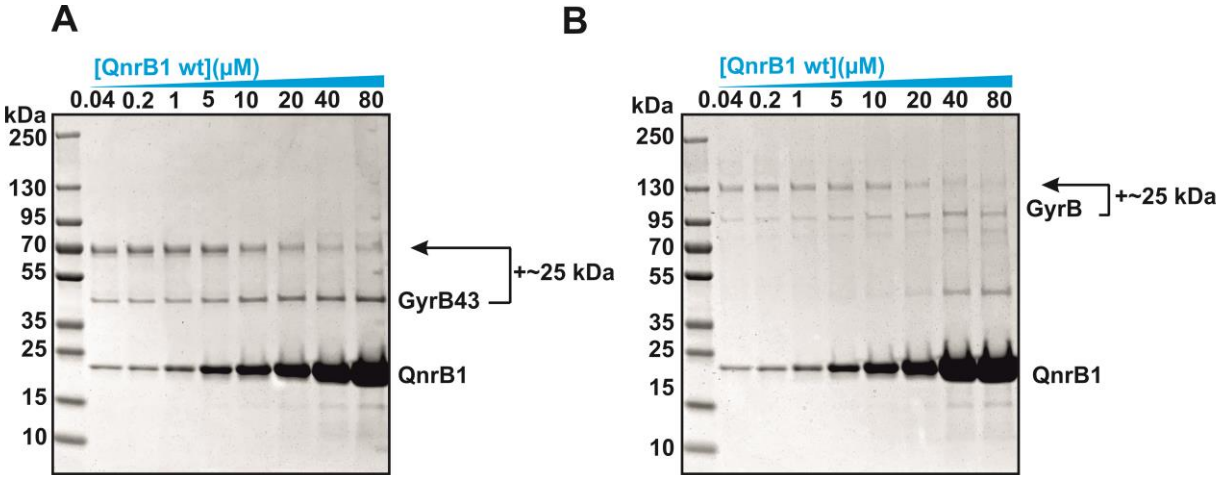


**Supplementary Figure S14. Competition crosslinking experiment with QnrB1 wt and QnrB1 Y123*p*Bpa.** (**A**) GyrB43 subunit. Each reaction consisted of 0.4 µM GyrB43, 5 µM QnrB1 Y123*p*Bpa and increasing concentration of WT unlabelled QnrB1 **(B)** GyrB subunit. Each reaction consisted of 0.4 µM GyrB, 5 µM QnrB1 Y123*p*Bpa and increasing concentrations of WT unlabelled QnrB1. Band-shifts corresponding to QnrB1 crosslinking are indicated. Coomassie-stained SDS-PAGE gels are shown.

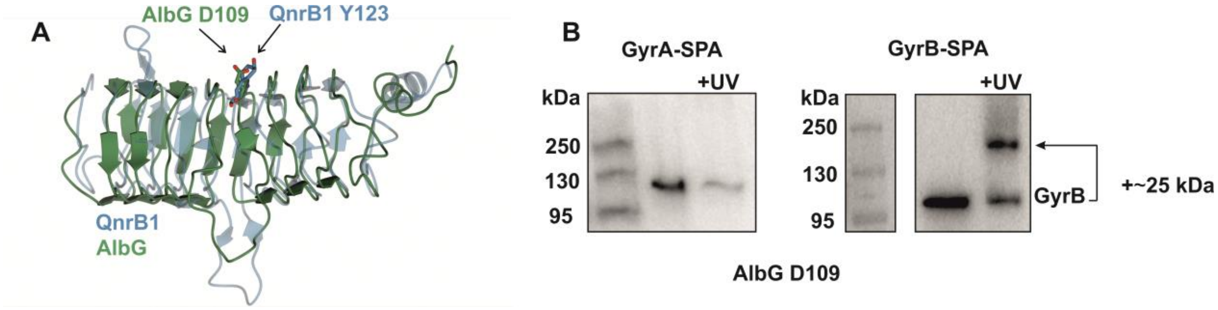


**Supplementary Figure S15. Structural alignment of QnrB1 and AlbG**. (**A**) Cartoon representations of QnrB1 (PDB:2XTW; blue) and AlbG (PDB:2XT2; green) superimposed by secondary structure matching in CCP4MG (1). Highlighted are QnrB1 Y123 and structurally homologous AlbG D109. (**B**). Western blot showing *in vivo* crosslinking of AlbG D109*p*Bpa to *E. coli* GyrA-SPA and GyrB-SPA.

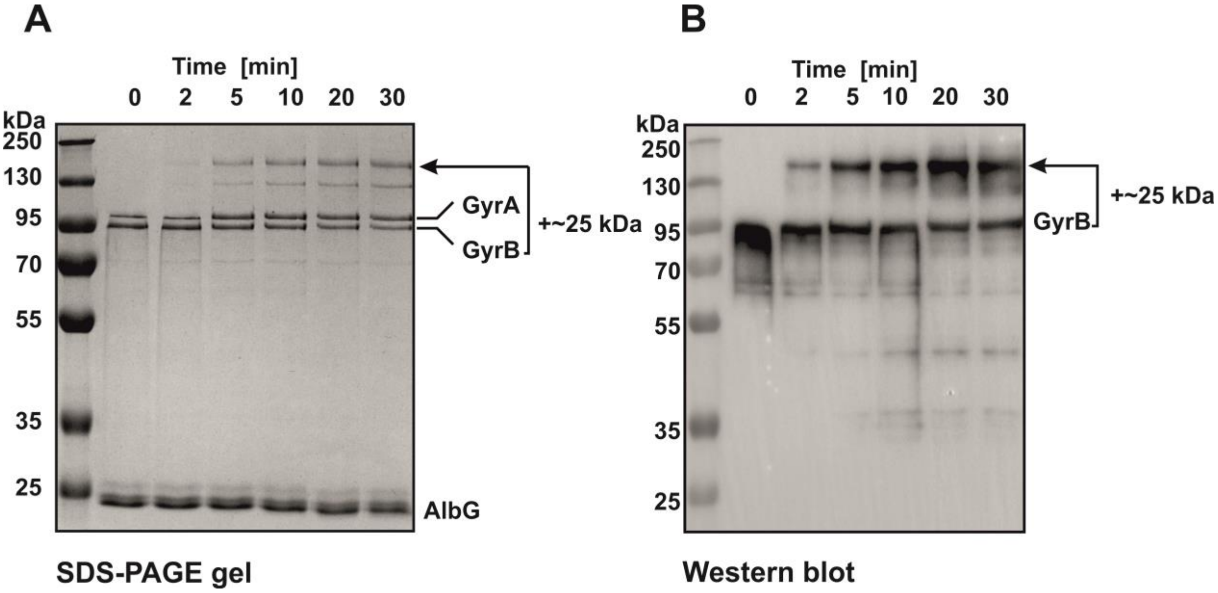


**Supplementary Figure S16. Time course of *in vitro* crosslinking of AlbG D109*p*Bpa to *E. coli* gyrase complex.** Each reaction included 0.4 μM gyrase complex (3xFLAG-tagged GyrB and untagged GyrA) and 5 μM AlbG D109*p*Bpa. (**A**) Coomassie-stained SDS-PAGE gel. (**B**) Western blot (α-FLAG). Band-shifts corresponding to QnrB1 crosslinking are indicated.

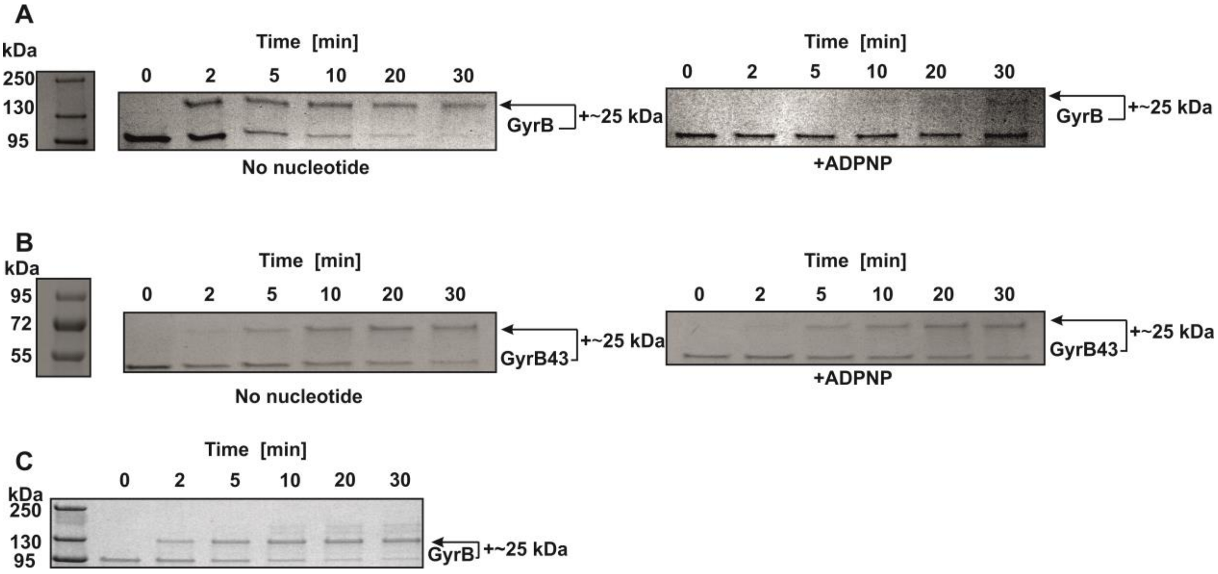


**Supplementary Figure S17. Time course of *in vitro* crosslinking of GyrB and GyrB43 to QnrB1Y123pBpa.** (**A**) Crosslink to gyrase B subunit with (*right*) and without (*left*) ADPNP. (**B**) Crosslink to gyrase B43 with (*right*) and without (*left*) ADPNP. (**C**) Crosslinking of QnrB1 Y123*p*Bpa ΔTTR mutant. Coomassie-stained SDS-PAGE gels are shown. Band-shifts corresponding to QnrB1 crosslinking are indicated.

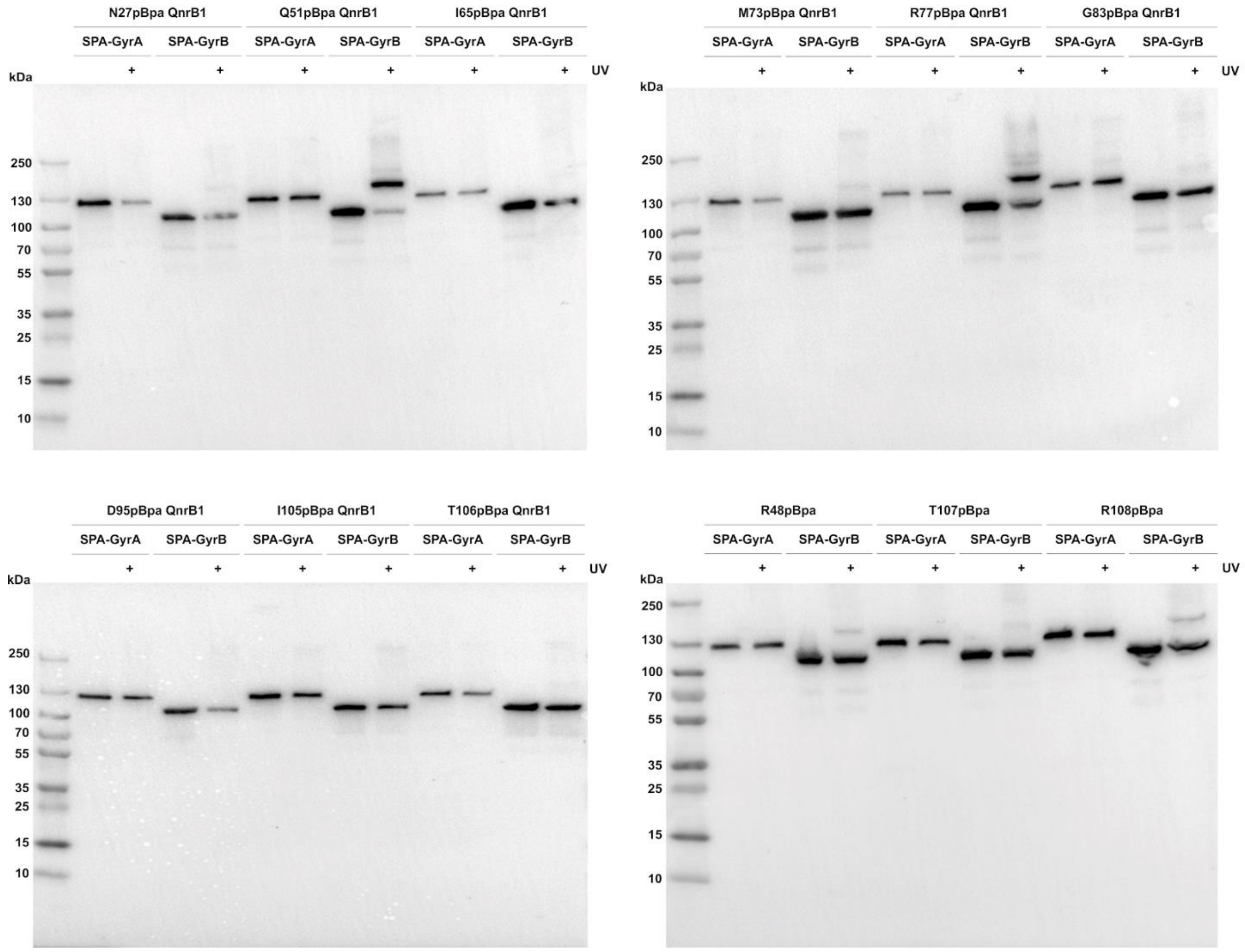


**Supplementary Figure S18. Uncropped Western blots for *in vivo* crosslinking of QnrB1 mutants to GyrA-SPA and GyrB-SPA.** Residues N27, Q51, I65, M73, R77, G83, D95, I105, T106, R48, T107, R108 are substituted to pBpa. Lanes with UV-treated cells are indicated by (+).

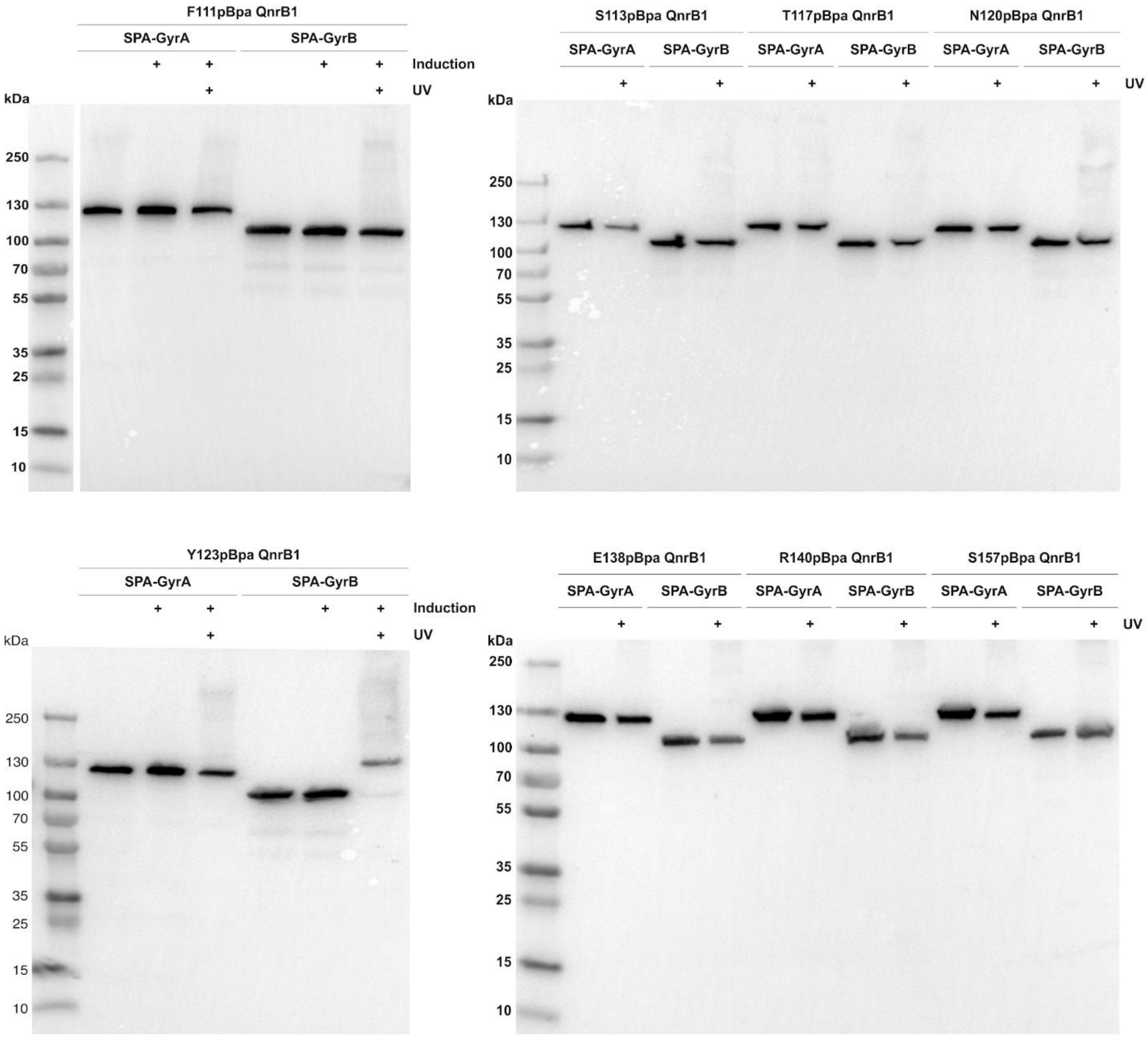


**Supplementary Figure S19. Uncropped Western blots for *in vivo* crosslinking of QnrB1 mutants to GyrA-SPA and GyrB-SPA.** Residues F111, S113, T117, N120, Y123, E138, R140, S157 are substituted to pBpa. Lanes with UV-treated cells are indicated by (+).

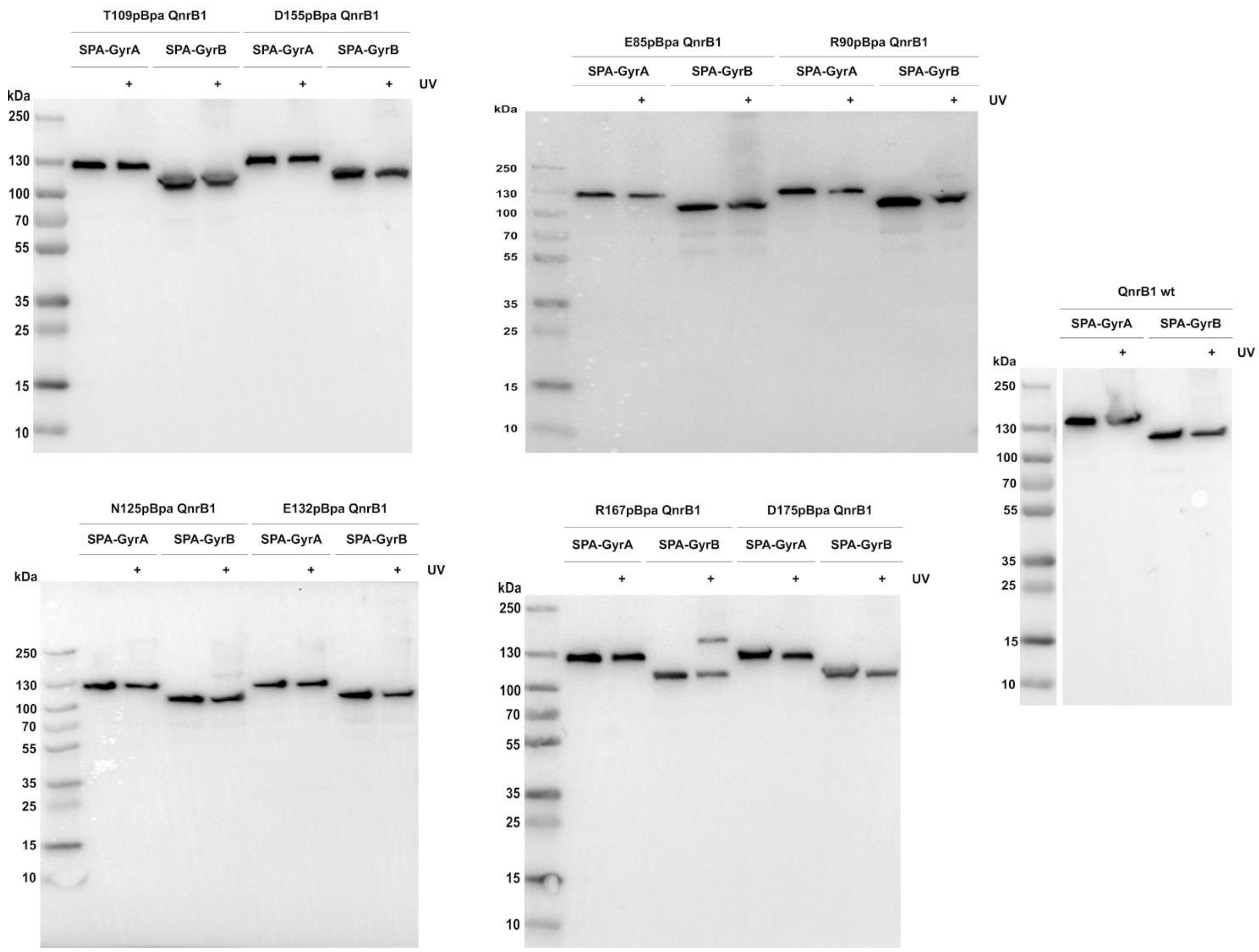


**Supplementary Figure S20. Uncropped Western blots for in vivo crosslinking of QnrB1 mutants to GyrA-SPA and GyrB-SPA.** Residues T109, D155, E85, R90, N125, E132, R167, D175 substituted to pBpa. Lanes with UV-treated cells are indicated by (+). Also shown is a negative control crosslinking with WT (unlabelled) QnrB1.

**Supplementary Table S1**. List of QnrB1 residues replaced with *p*Bpa; **bold** are residues, which produced UV crosslinks

| **QnrB1 position** | **PRP position** (2) |
|---|---|
| N27 | $i^{+2}$ |
| R48 | loop A |
| **Q51** | loop A |
| I65 | $i^{-1}$ |
| M73 | $i^{+2}$ |
| **R77** | $i^{+1}$ |
| G83 | $i^{+2}$ |
| E85 | $i^{-1}$ |
| R90 | $i^{-1}$ |
| D95 | $i^{-1}$ |
| I105 | loop B |
| T106 | loop B |
| T107 | loop B |
| R108 | loop B |
| T109 | loop B |
| F111 | loop B |
| S113 | loop B |
| T117 | $i^{+1}$ |
| N120 | $i^{-1}$ |
| **Y123** | $i^{+1}$ |
| N125 | $i^{-1}$ |
| E132 | $i^{+1}$ |
| E138 | $i^{+2}$ |
| R140 | $i^{-1}$ |
| D155 | $i^{-1}$ |
| S157 | $i^{+1}$ |
| **R167** | $i^{+1}$ |
| D175 | $i^{-1}$ |
| R187 | $i^{+1}$ |

**Table S2. List of plasmids used in the study.**

| Name | Backbone | Source | Purpose |
|---|---|---|---|
| pBAD-*mcbG* | pBAD/His B | this work | Expression of untagged McbG protein |
| pBAD-*albG* | pBAD/His B | this work | Expression of untagged AlbG protein |
| pBAD-*albG* Δ91–97 | pBAD/His B | this work | Expression of untagged AlbG loop deletion mutant |
| pBAD-*qnrb1* | pBAD/His B | this work | Expression of untagged QnrB1 protein |
| pET28- *mcbG* | pET-28a(+) | this work | Expression of His tagged McbG protein |
| pET28-*albG* | pET-28a(+) | Gift of Dr. Mikhail Metelev (Uppsala University) | Purification of 6xHis-tagged AlbG protein |
| pET28- *qnrb1* | pET-28a(+) | Gift of Dr. Mikhail Metelev (Uppsala University) | Purification of 6xHis-tagged QnrB1 protein |
| pBAD-*mcbABCDEFG* | pBAD/His B | Gift of Dr. Mikhail Metelev (Uppsala University) | Encodes microcin B biosynthetic cluster (microcin B production) |
| pET21-GyrA | pET-21b(+) | this work (Dr. Jonathan Heddle) | Purification of untagged *E.coli* GyrA |
| pET21-GyrB | pET-21b(+) | this work (Dr. Jonathan Heddle) | Purification of untagged *E. coli* GyrB |
| pET21-3xFLAG-GyrB | pET-21b(+) | this work | Purification of FLAG-tagged GyrB |
| pET21-GyrA-FLAG | pET-21b(+) | this work | Purification of FLAG-tagged GyrA |
| pET28-GyrB47 | pET-28a(+) | this work | Purification of 6xHis-GyrB47 |
| pAJR10.18 (GyrB47) | pET-21a(+) | Gift of Anthony. Maxwell, (John Innes Centre) | Purification of GyrB47 (untagged) |
| pAJ1 (GyrB43) | pET-21a(+) | Gift of Anthony. Maxwell, (John Innes Centre) | Purification of GyrB43 (untagged) |
| pLIC172-escoGyrBAfus | pLIC | Gift of James Berger, (John Hopkins School of Medicine) | Purification of *E. coli* GyrBAcore protein |
| pET28- HIS_FLAG_*qnrb1* | pET-28a(+) | this work | Purification of FLAG- and 6xHis-tagged QnrB1 |
| pET28- HIS_FLAG_*albG* | pET-28a(+) | this work | Purification of FLAG- and 6xHis-tagged AlbG |
| pET28- HIS_*qnrb1ΔTTR* | pET-28a(+) | this work | Purification of QnrB1 loop deletion mutant |
| pEVOL-pBpF | pEVOL | Addgene #31190 | Incorporation of *p*Bpa |
| pBAD-HIS_*qnrb1*_pBpa | pBAD/His B | this work | Incorporation of *p*Bpa |
| pBAD-HIS_*albG*_pBpa | pBAD/His B | this work | Incorporation of *p*Bpa |

### Table S3. List of primers used in the study

| Primer Name | Primer sequence (5'-3') | Purpose |
|---|---|---|
| 300f | CGGTATTCGGAATCTTGCAC | Amplification of pBR322 300 bp fragment |
| 300r | GCGGTCCAATGATCGAAG | Amplification of pBR322 300 bp fragment |
| 220f | CACTGGTCCCGCCACC | Amplification of pBR322 220 bp fragment |
| 220r | CGATCCTTGAAGCTGTCC | Amplification of pBR322 220 bp fragment |
| 147f | AGGCCATTATCGCCGGCATG | Amplification of pBR322 147bp fragment |
| 147r | GCCTGGACAGCATGGCCTG | Amplification of pBR322 147bp fragment |
| 133f | TATCGCCGGCATGGCGGC | Amplification of pBR322 133 bp fragment |
| 133r | CAGCATGGCCTGCAACGC | Amplification of pBR322 133 bp fragment |
| 100f | CGACGCGCTGGGCTACGTC | Amplification of pBR322 100 bp fragment |
| 100r | CGCGGGCATCCCGATGCCG | Amplification of pBR322 100 bp fragment |
| 76f | CTACGTCTTGCTGGCGTTCGCGACGCGAGGCTGGATGGCCTTCCCCA TTATGATTCTTCTCGCTTCCGGCGGCATC | Annealing of 76 bp pBR322 fragment |
| 76r | GATGCCGCCGGAAGCGAGAAGAATCATAATGGGGAAGGCCATCCA GCCTCGCGTCGCGAACGCCAGCAAGACGTAG | Annealing of 76 bp pBR322 fragment |
| For_NcoI_McbG | ATCCCATGGGGATGGATATAATAGAAAAAAGAATCACAAAACGA | Cloning of tagless *mcbG* gene into pBAD HIS B plasmid |
| Rev_XhoI_McbG | ATCTCTCGAGTCATCCCCCTACAACCACTC | |
| For_NcoI_AlbG | ATCCCATGGGGATGCCGGCCAAGACCCTTGAAAGCAAGG | Cloning of tagless *albG* gene into pBAD HIS B plasmid |
| Rev_XhoI_Albg | ATCTCTCGAGTCAATCGGACAGCTCGATATCCAGGCT | |
| For_ AlbG_Δ91–97 | GTCAACTGGACCAGCGCACAAGCGGGGGCGCTGTCGTTCGAGCGCT G | Cloning of tagless *albG* loop deletion Δ91–97 gene into pBAD HIS B plasmid |
| Rev_AlbG_Δ91–97 | CAGCGCTCGAACGACAGCGCCCCCGCTTGTGCGCTGGTCCAGTTGA C | Cloning of tagless *albG* loop deletion Δ91–97 gene into pBAD HIS B plasmid |
| For_NcoI_QnrB1 | ATCCCATGGGGATGGCTCTGGCACTCGTTGGCGAAA | Cloning of tagless *qnrb1* gene into pBAD HIS B plasmid |

| | | |
|---|---|---|
| Rev_XhoI_QnrB1 | CCGCTCGAGTTAACCAATCACCGCGATGCCAAGTCGCTCCAT | Cloning of tagless *qnrb1* gene into pBAD HIS B plasmid |
| For_NdeI_3xFLAG-GyrB | AATACATATGGACTACAAAGACCATGACGGTGATTATAAAGATCATGACATCGATTACAAGGATGACGATGACAAGTCGAATTCTTATGACTCCTC | Cloning of N-terminally 3xFLAG *E .coli gyrB* gene |
| Rev_GyrB_XhoI | AATACTCGAGTTAAATATCGATATTCGCCGCTTTCAGG | Cloning of *E. coli gyrB* gene |
| For_NdeI_GyrA | TTATCATATGAGCGACCTTGCGAGAG | Cloning of *E. coli gyrA* gene |
| Rev_XhoI_GyrA_FLAG | TTAACTCGAGTTACTTGTCGTCATCGTCTTTGTAGTCACCGCTACCTTCTTCTTCTGGCTCGTCGTC | Cloning of C-terminally FLAG-tagged *E. coli gyrA* gene |
| For_NdeI_GyrB47 | AATACATATGCGCCGTAAAGGTGCGC | Cloning of *E. coli* GyrB47 (TOPRIM) subdomain |
| For_NcoI_6xHis_FLAG_QnrB1 | ATTACCATGGGCCATCATCATCATCATCATAGCGGCGATTATAAGGACGATGACGATAAGAGCGGCATGGCTCTGGCACTCGTTGGCGAAA | Cloning of N-terminally 6xHIS FLAG *qnrB1* gene |
| For_NcoI_6xHis_FLAG_AlbG | ATTACCATGGGCCATCATCATCATCATCATAGCGGCGATTATAAGGACGATGACGATAAGAGCGGCATGCCGGCCAAGACCCTTG | Cloning of N-terminally 6xHIS FLAG *albG* gene |
| For_AlbG_D109TAG | TGCATCCTCAACTAGAGCTTGTTCTAC | Introducing amber stop codon in *albG* gene |
| Rev_AlbG_D109TAG | GTAGAACAAGCTCTAGTTGAGGATGCA | Introducing amber stop codon in *albG* gene |
| For_Qnrb1_N27TAG | ACATTTTTTTAGTGTGATTTTTCA | Introducing amber stop codon in *qnrb1* gene |
| Rev_Qnrb1_N27TAG | TGAAAAATCACACTAAAAAAATGT | − |
| For_Qnrb1_R48TAG | CAGTTCTATGATTAGGAAAGCCAGAAA | − |
| Rev_Qnrb1_R48TAG | TTTCTGGCTTTCCTAATCATAGAACTG | − |
| For_Qnrb1_Q51TAG | GATCGTGAAAGCTAGAAAGGGTGC | − |
| Rev_Qnrb1_Q51TAG | GCACCCTTTCTAGCTTTCACGATC | − |
| For_Qnrb1_I65TAG | CTGAAAGATGCCTAGTTTAAAAGC | − |
| Rev_Qnrb1_I65TAG | GCTTTTAAACTAGGCATCTTTCAG | − |
| For_Qnrb1_M73TAG | AGCTGTGATTTATCATAGGCGGATTTT | − |
| Rev_Qnrb1_M73TAG | AAAATCCGCCTATGATAAATCACAGCT | − |
| For_Qnrb1_R77TAG | GCGGATTTTTAGAATTCCAGTGCG | − |
| Rev_Qnrb1_R77TAG | CGCACTGGAATTCTAAAAATCCGC | − |
| For_Qnrb1_N78TAG | GCGGATTTTCGCTAGTCCAGTGCGCTG | − |
| Rev_Qnrb1_N78TAG | CAGCGCACTGGACTAGCGAAAATCCGC | − |
| For_Qnrb1_G83TAG | AGTGCGCTGTAGATTGAAATT | − |
| Rev_Qnrb1_G83TAG | AATTTCAATCTACAGCGCACT | − |
| For_Qnrb1_E85TAG | CTGGGCATTTAGATTCGCCAC | − |
| Rev_Qnrb1_E85TAG | GTGGCGAATCTAAATGCCCAG | − |
| For_Qnrb1_R90TAG | TGAAATTCGCCACTGCTAGGCACAAGG | − |
| Rev_Qnrb1_R90TAG | CCTTGTGCCTAGCAGTGGCGAATTTCA | − |
| For_Qnrb1_D95TAG | ACAAGGCGCATAGTTCCGCGGC | − |
| Rev_Qnrb1_D95TAG | GCCGCGGAACTATGCGCCTTGT | − |
| For_Qnrb1_I105TAG | ATGAATATGTAGACCACGCGCACC | − |
| Rev_Qnrb1_I105TAG | GGTGCGCGTGGTCTACATATTCAT | − |
| For_Qnrb1_T106TAG | AATATGATCTAGACGCGCACCTGG | − |
| Rev_Qnrb1_T106TAG | CCAGGTGCGCGTCTAGATCATATT | − |
| For_Qnrb1_T107TAG | AATATGATCACCTAGCGCACCTGGTTT | − |
| Rev_Qnrb1_T107TAG | AAACCAGGTGCGCTAGGTGATCATATT | − |

| | | |
|---|---|---|
| For_Qnrb1_R108TAG | ATGATCACCACGTAGACCTGGTTTTGT | – |
| Rev_Qnrb1_R108TAG | ACAAAACCAGGTCTACGTGGTGATCAT | – |
| For_Qnrb1_T109TAG | ATCACCACGCGCTAGTGGTTTTGTA | – |
| Rev_Qnrb1_T109TAG | TACAAAACCACTAGCGCGTGGTGAT | – |
| For_Qnrb1_F111TAG | CGCACCTGGTAGTGTAGCGCATAT | – |
| Rev_Qnrb1_F111TAG | ATATGCGCTACACTACCAGGTGCG | – |
| For_Qnrb1_S113TAG | TGGTTTTGTTAGGCATATATCACG | – |
| Rev_Qnrb1_S113TAG | CGTGATATATGCCTAACAAAACCA | – |
| For_Qnrb1_T117TAG | AGCGCATATATCTAGAATACCAATCTA | – |
| Rev_Qnrb1_T117TAG | TAGATTGGTATTCTAGATATATGCGCT | – |
| For_Qnrb1_N120TAG | ACGAATACCTAGCTAAGCTACGCC | – |
| Rev_Qnrb1_N120TAG | GGCGTAGCTTAGCTAGGTATTCGT | – |
| For_Qnrb1_Y123TAG | ACCAATCTAAGCTAGGCCAATTTTTCG | – |
| Rev_Qnrb1_Y123TAG | CGAAAAATTGGCCTAGCTTAGATTGGT | – |
| For_Qnrb1_N125TAG | AGCTACGCCTAGTTTTCGAAAGTC | – |
| Rev_Qnrb1_N125TAG | GACTTTCGAAAACTAGGCGTAGCT | – |
| For_Qnrb1_E132TAG | GTCGTGTTGTAGAAGTGTGAGCTG | – |
| Rev_Qnrb1_E132TAG | CAGCTCACACTTCTACAACACGAC | – |
| For_Qnrb1_E138TAG | TGTGAGCTGTGGTAGAACCGTTGG | – |
| Rev_Qnrb1_E138TAG | CCAACGGTTCTACCACAGCTCACA | – |
| For_Qnrb1_R140TAG | CTGTGGGAAAACTAGTGGATAGGT | – |
| Rev_Qnrb1_R140TAG | ACCTATCCACTAGTTTTCCCACAG | – |
| For_Qnrb1_L147TAG | GGTGCCCAGGTATAGGGCGCGACGTTC | – |
| Rev_Qnrb1_L147TAG | GAACGTCGCGCCCTATACCTGGGCACC | – |
| For_Qnrb1_D155TAG | TTCAGTGGTTCATAGCTCTCC | – |
| Rev_Qnrb1_D155TAG | GGAGAGCTATGAACCACTGAA | – |
| For_Qnrb1_S157TAG | TTCAGATCTCTAGGGCGGCGA | – |
| Rev_Qnrb1_S157TAG | TCGCCGCCCTAGAGATCTGAA | – |
| For_Qnrb1_R167TAG | ACTTTCGACTGGTAGGCAGCAAAC | – |
| Rev_Qnrb1_R167TAG | GTTTGCTGCCTACCAGTCGAAAGT | – |
| For_Qnrb1_D175TAG | ACACATTGCTAGCTGACCAATTCG | – |
| Rev_Qnrb1_D175TAG | CGAATTGGTCAGCTAGCAATGTGT | – |
| For_Qnrb1_D185TAG | TTGGGTGACTTATAGATTCGGGGC | – |
| Rev_Qnrb1_D185TAG | GCCCCGAATCTATAAGTCACCCAA | – |
| For_Qnrb1_R187TAG | GACTTAGATATTTAGGGCGTTGAT | – |
| Rev_Qnrb1_R187TAG | ATCAACGCCCTAAATATCTAAGTC | – |
